# Pan-cancer nucleosome landscapes reveal tumor heterogeneity and inform cell-free DNA-based detection

**DOI:** 10.64898/2026.10.12.758417

**Authors:** Shen Yang, Jialuo Ye, Jinqi Yuan, Xiaoqian Huang, Yue Yin, Zhaoying Liu, Wei Li, Yumei Li

## Abstract

**Background:** Nucleosome positioning and occupancy are fundamental features of chromatin organization, yet their variation across human cancers and potential utility for noninvasive cancer detection remain poorly characterized.

**Methods:** We systematically characterized nucleosome organization across 410 primary tumors spanning 23 cancer types using ATAC-seq, establishing a high-confidence pan-cancer nucleosome atlas validated by independent MNase-seq data. We characterized shared and cancer-type-enriched nucleosome patterns, investigated their associations with tumor heterogeneity and regulatory alterations, and evaluated tissue-derived nucleosome signatures for cancer classification and detection using circulating cell-free DNA (cfDNA).

**Results:** The atlas identified shared and cancer-type-enriched nucleosomes associated with distinct regulatory programs. Notably, nucleosome occupancy provided information complementary to chromatin accessibility for tumor stratification and revealed molecular heterogeneity within individual cancers, including subgroups associated with Basal-like breast cancer. In hepatocellular carcinoma, differential nucleosome analysis revealed cancer-associated chromatin remodeling involving regulatory pathways, transcription factor motifs, and candidate genes. Importantly, tissue-derived nucleosome signatures were detectable in cfDNA, enabling classification among three cancer types (AUROC = 0.877) and colorectal cancer detection (AUROC = 0.930), with an AUROC of 0.773 in an independent validation cohort. We also developed OncoNucAtlas, a publicly accessible database facilitating exploration of these nucleosome landscapes.

**Conclusions:** Our study cancer-associated nucleosome organization as a complementary source of epigenomic information and demonstrates the potential of tissue-derived nucleosome signatures for cfDNA-based cancer detection. The pan-cancer atlas provides a resource for investigating tumor heterogeneity and developing noninvasive cancer detection approaches.

## Introduction

In eukaryotic cells, genomic DNA is packaged into nucleosomes, the fundamental units of chromatin, each consisting of approximately 147 bp of DNA wrapped around a histone octamer^1^. The *in vivo* nucleosome organization is not random but is shaped by both intrinsic DNA sequence preferences^2^ and trans-acting factors, such as chromatin remodelers and transcription factors (TFs)^3^. Well-defined nucleosome patterns, such as phased nucleosome arrays, are frequently observed around promoters^4,5^ and TF binding sites (e.g. CTCF^6^). On the other hand, nucleosome positioning can modulate TF binding by controlling the accessibility of regulatory elements^7^. Therefore, nucleosome organization is closely linked to gene regulation^8,9^ and dynamically responds to cellular stimuli and pathological perturbations^10^.

Especially, nucleosome organization is extensively altered in cancer, with important consequences for oncogenic gene regulation. For example, nucleosome gain at enhancers can promote DNA hypermethylation and gene silencing, whereas nucleosome loss is often associated with increased chromatin accessibility and aberrant gene activation^11^. Cancer-associated alterations in chromatin remodeling machinery can further disrupt nucleosome organization. Mutations in chromatin remodeling complexes, such as the SWI/SNF complex, can perturb nucleosome positioning at regulatory regions and reshape transcriptional programs that contribute to malignant phenotypes^12^. These observations suggest that nucleosome organization represents an important layer of epigenetic regulation through which cancer cells acquire and maintain aberrant regulatory states.

Importantly, these cancer-associated nucleosome alterations can also leave measurable signatures in circulating cell-free DNA (cfDNA), providing a potential link between tissue chromatin states and non-invasive cancer detection. Because cfDNA fragmentation is influenced by nucleosome protection and chromatin accessibility in its cells of origin, features such as nucleosome footprints^13^ and fragment size distributions^14^ retain information about the underlying chromatin architecture. Notably, cfDNA derived from tumor cells exhibits distinct fragmentomic alterations, such as an increased abundance of shorter fragments and altered fragmentation patterns around regulatory regions, which are associated with the underlying nucleosome organization and chromatin accessibility^15–18^. These properties have motivated the use of nucleosome-associated cfDNA features for cancer detection and tissue-of-origin inference. However, despite the availability of genome-wide nucleosome maps in humans, most existing studies have focused on specific regulatory regions or individual cancer types^19–21^. A systematic reference of nucleosome organization across diverse human cancers, and its potential for identifying cancer-associated regulatory alterations and informing cfDNA analysis, remains lacking.

Currently, micrococcal nuclease digestion with deep sequencing (MNase-seq^22^), which preferentially digests linker DNA between nucleosomes, remains a widely used approach for generating high-resolution, genome-wide nucleosome maps. However, generating MNase-seq profiles across large numbers of primary tumor samples remains challenging, limiting its application to systematic pan-cancer analyses. In contrast, the assay for transposase-accessible chromatin using sequencing (ATAC-seq^23^) has been extensively applied to primary tumors and provides information not only on chromatin accessibility but also on nucleosome organization through the characteristic distribution of nucleosome-protected fragments. Computational methods such as NucleoATAC^24^ and deNOPA^25^ can leverage these signals to infer nucleosome positions from ATAC-seq data. The growing availability of large-scale cancer ATAC-seq datasets therefore provides an opportunity to systematically characterize nucleosome organization across tumor types and investigate regulatory information beyond conventional accessibility-based analyses.

Thus, we constructed a pan-cancer nucleosome atlas from ATAC-seq profiles of primary tumors across 23 human cancer types in The Cancer Genome Atlas (TCGA). We systematically characterized shared and cancer-type-enriched patterns of nucleosome organization and examined their relationships with chromatin accessibility, regulatory programs, and tumor heterogeneity. We further investigated nucleosome-defined molecular subtypes and evaluated whether tissue-derived nucleosome signatures are retained in cfDNA and can inform cancer classification and detection. Together, our study provides a systematic view of nucleosome organization across human cancers and establishes a framework for investigating its biological and translational relevance.

## Results

### A robust pan-cancer nucleosome atlas derived from ATAC-seq

To systemically characterize the pan-cancer nucleosome landscapes, we first established a strategy for inferring nucleosome positions from ATAC-seq data by defining high-confidence nucleosome positions as the overlap between predictions generated by two dedicated algorithms, deNOPA^25^ and NucleoATAC^24^. Because deNOPA and NucleoATAC rely on distinct statistical models for nucleosome inference, only nucleosomes supported by both methods were retained to improve robustness and reduce caller-specific bias. We validated this strategy using four datasets^26–29^ with matched ATAC-seq and MNase-seq data (**Table S1**). Notably, a substantial proportion of nucleosomes inferred from ATAC-seq overlapped with those identified by MNase-seq (**Fig. S1A**), and their occupancy profiles were highly concordant between the two assays (**Fig. S1B**). Moreover, nucleosome positions inferred from ATAC-seq exhibited well-phased and consistent nucleosome signals in both ATAC-seq and MNase-seq data (**Fig. S2**). Together, these results demonstrate that ATAC-seq can reliably recover nucleosome positioning and occupancy.

Having established a reliable nucleosome inference strategy, we next applied it to ATAC-seq profiles from 410 primary tumors spanning 23 cancer types in The Cancer Genome Atlas (TCGA)^30^. By identifying reproducible nucleosome positions (detected in at least two samples) within each cancer type, we constructed a high-confidence pan-cancer nucleosome atlas (**Fig. S3; Methods**). On average, each cancer type contained 918,901 nucleosome positions (range, 121,043-2,385,544), spanning both regulatory elements and distal intergenic regions (**Fig. 1A**). Consistent with canonical chromatin architecture, inferred nucleosomes exhibited well-phased positioning around CTCF binding sites and the characteristic nucleosome-depleted region flanked by regularly positioned nucleosomes at transcription start sites (TSSs) (**Fig. 1B**), in agreement with previous reports^31,32^.

**Fig. 1.**
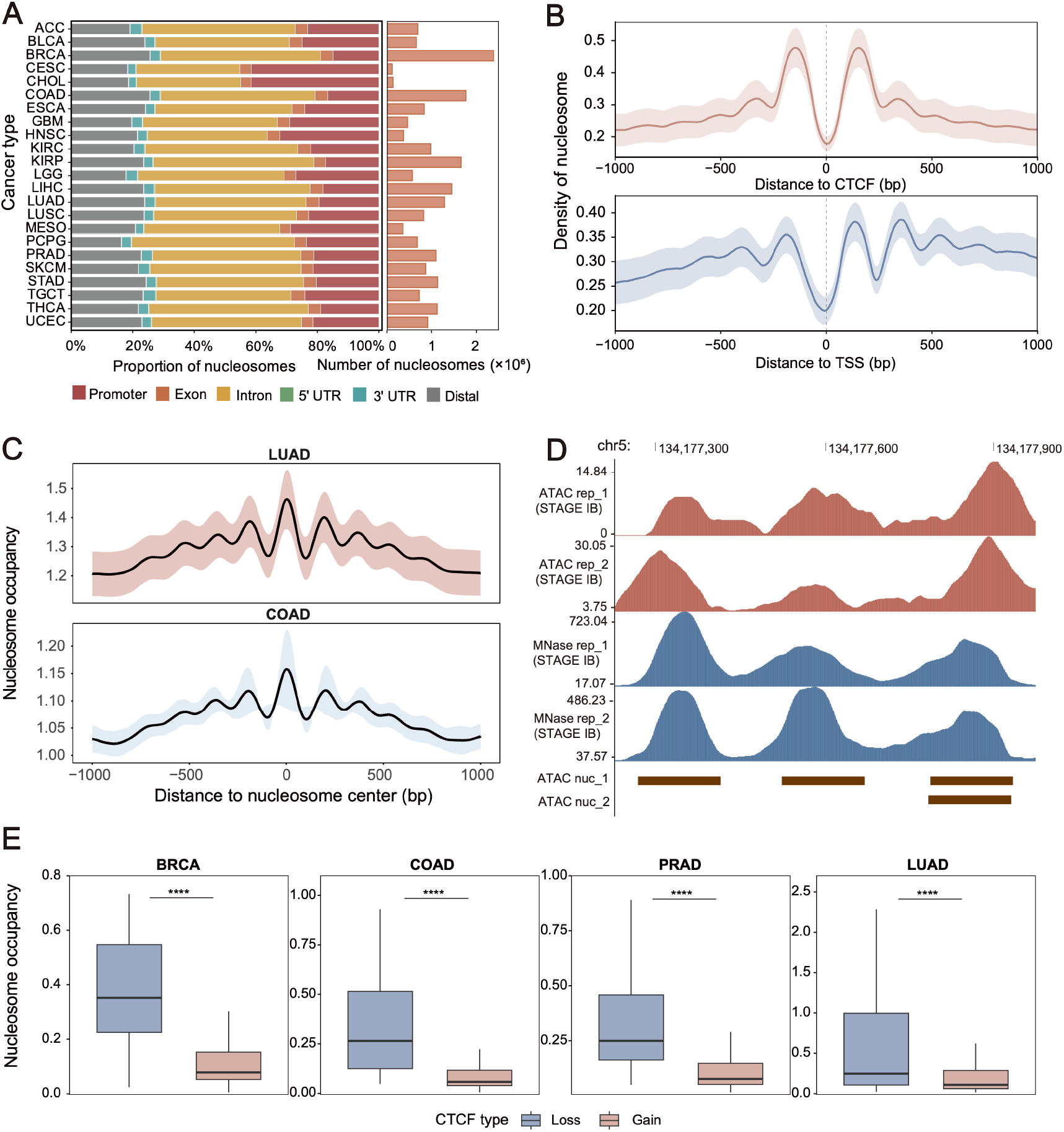
Characteristics and validation of the pan-cancer nucleosome landscape inferred from ATAC-seq. **(A)** Genomic distribution and number of cancer-type-level nucleosomes across the 23 cancer types included in the pan-cancer nucleosome atlas. **(B)** Aggregative nucleosome density profiles around transcription start sites (TSSs) and constitutive CTCF binding sites for cancer-type-level nucleosomes. Lines represent the mean profile across all 23 cancer types, with the shaded area representing 95% confidence intervals. **(C)** Distribution of MNase-seq nucleosome occupancy around ATAC-seq-inferred nucleosome positions in LUAD and COAD. Lines represent the mean profile across samples, with the shaded area representing 95% confidence intervals. **(D)** Representative genomic regions showing nucleosome occupancy signals derived from ATAC-seq (red) and MNase-seq (blue). Nucleosome positions inferred from ATAC-seq are shown below the occupancy tracks. ATAC-seq data were obtained from TCGA LUAD samples, and MNase-seq data were obtained from an independent public LUAD dataset (GSE74340) with matched tumor stage. **(E)** Nucleosome occupancy from ATAC-seq data at CTCF-binding sites showing gain or loss of CTCF occupancy in tumors relative to normal tissues across four cancer types. Gain and loss CTCF sites were defined based on differential CTCF binding between tumor and normal samples as described in Zang *et al.*^34^. Differences in nucleosome occupancy between CTCF gain and loss sites were assessed using a two-sided Wilcoxon rank-sum test. Statistical significance between groups is denoted by asterisks (****P < 0.0001).

To further validate the inferred nucleosome positions, we analyzed the only two publicly available tumor-derived MNase-seq datasets matching the cancer types examined in our study, generated from lung adenocarcinoma (LUAD) and colon adenocarcinoma (COAD) tissues^33^. Nucleosome occupancy profiles derived from MNase-seq displayed clear phasing and periodicity centered on the ATAC-seq-inferred nucleosome positions (**Fig. 1C**). At representative genomic loci, nucleosome occupancy patterns inferred from ATAC-seq closely matched those measured by MNase-seq in tumors of the same cancer type (**Fig. 1D**), providing independent validation of the accuracy and robustness of our pan-cancer nucleosome atlas.

To investigate whether the atlas captures cancer-specific chromatin organization, we further examined nucleosome patterns surrounding cancer-associated CTCF binding sites from a previous study^34^, which define cancer-specific CTCF gain and loss sites relative to normal samples across four cancer types. Compared with CTCF loss sites, gain sites displayed lower nucleosome occupancy in tumors (**Fig. 1E**), consistent with increased chromatin accessibility accompanying CTCF binding. These results demonstrate that our pan-cancer nucleosome atlas faithfully recapitulates both canonical nucleosome organization and cancer-associated chromatin remodeling.

### Shared and cancer-type-enriched nucleosome landscapes across human cancers

Although the pan-cancer nucleosome atlas may reveal shared nucleosome organization across human cancers, we reasoned that substantial epigenetic heterogeneity should also exist in the form of cancer-type-enriched nucleosome positioning. Building upon the pan-cancer nucleosome atlas, we therefore classified nucleosomes into pan-cancer nucleosomes (**Table S2**), consistently observed across all cancer types, and cancer-type-enriched nucleosomes, exhibiting high occupancy in individual cancer types (**Fig. S3; Methods**). To independently validate these cancer-type-enriched nucleosomes, we analyzed the two publicly available tumor-derived MNase-seq datasets from COAD and LUAD. In both cancer types, nucleosome occupancy measured by MNase-seq was significantly higher at the corresponding ATAC-seq-derived cancer-type-enriched nucleosome positions (**Fig. 2A**), providing orthogonal evidence that these inferred nucleosome landscapes faithfully represent bona fide chromatin organization.

**Fig. 2.**
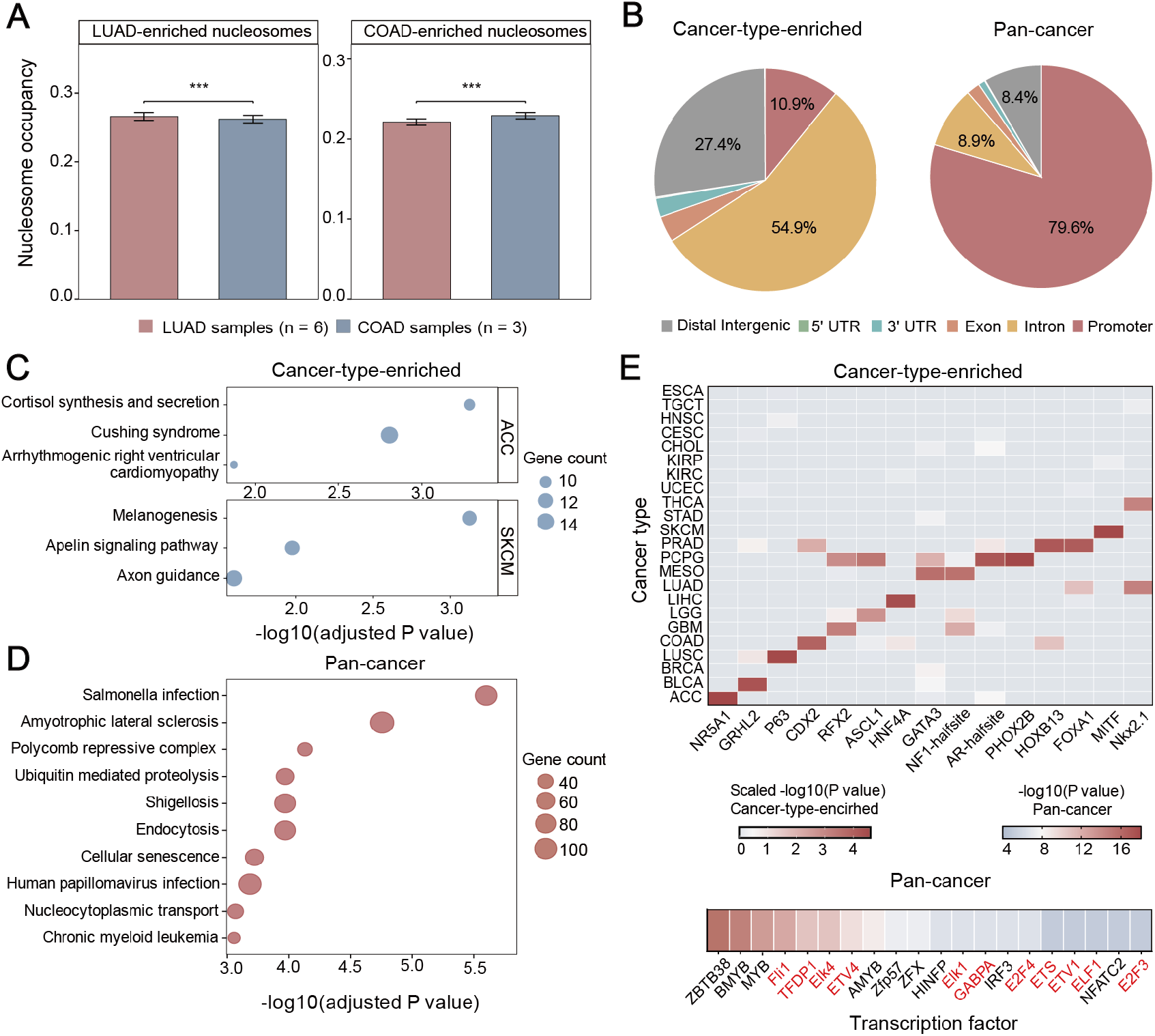
Cancer-type-enriched and pan-cancer nucleosomes reveal distinct genomic and functional characteristics. **(A)** Nucleosome occupancy measured by MNase-seq (GSE74340) at LUAD- and COAD-enriched nucleosome regions. Bars represent the mean nucleosome occupancy, and error bars indicate the standard error of the mean (SEM). Statistical significance between groups is denoted by asterisks (\*\*\**P* < 0.001). **(B)** Genome distribution of pan-cancer nucleosomes and cancer-type-enriched nucleosomes. **(C)** Dot plots showing the top three enriched pathways for genes associated with the top 1,000 cancer-type-enriched nucleosomes in ACC and SKCM. **(D)** Dot plots showing the top 20 enriched pathways for genes associated with pan-cancer nucleosomes. **(E)** Heatmaps showing transcription factor (TF) motif enrichment in cancer-type-enriched and pan-cancer nucleosomes. For cancer-type-enriched nucleosomes, 15 lineage-associated TF motifs were evaluated across cancer types, and motif enrichment significance [−log10(P value)] was scaled across cancer types for each TF motif (column). For pan-cancer nucleosomes, the 20 most significantly enriched TF motifs are shown.

We next compared the genomic and functional characteristics of cancer-type-enriched and pan-cancer nucleosomes. Genomic annotation revealed distinct distributions, with cancer-type-enriched nucleosomes broadly distributed across genomic elements whereas pan-cancer nucleosomes were preferentially enriched at promoter regions (**Fig. 2B**). Functional enrichment analysis further revealed that cancer-type-enriched nucleosomes were associated with pathways reflecting tissue identity and cancer-specific biology (**Fig. 2C, Fig. S4, and Table S3**), including melanogenesis in skin cutaneous melanoma (SKCM)^35^, as well as cortisol synthesis and secretion in adrenocortical carcinoma (ACC)^36^. In contrast, pan-cancer nucleosomes were enriched in core cellular processes, including ubiquitin mediated proteolysis, endocytosis, and cellular senescence (**Fig. 2D**). Cancer-type-enriched nucleosomes were also significantly enriched for motifs of lineage-associated transcription factors^30^ (**Fig. 2E and Table S4**), including MITF in SKCM^37^, and TP63 in lung squamous cell carcinoma (LUSC)^38^, whereas pan-cancer nucleosomes preferentially enriched motifs of the ETS and E2F transcription factor families (**Fig. 2E and Table S5**). Together, these findings suggest that pan-cancer nucleosomes preferentially mark regulatory elements associated with fundamental cellular functions, whereas cancer-type-enriched nucleosomes capture lineage-dependent regulatory programs underlying tumor-specific chromatin organization.

### Cancer-type-enriched nucleosomes complement chromatin accessibility for tumor stratification

We next investigated whether cancer-type-enriched nucleosome positioning simply reflects chromatin accessibility or provides complementary information for tumor stratification. Only 28.90% of cancer-type-enriched nucleosomes overlapped with cancer-type-enriched ATAC-seq peaks, with overlap rates ranging from 4.21% to 17.34% across individual cancer types (**Fig. 3A**). Nucleosome positions overlapping cancer-type-enriched ATAC-seq peaks were predominantly associated with active regulatory processes (**Fig. 3B and Table S6**), consistent with coordinated changes in nucleosome organization and chromatin accessibility. In contrast, nucleosome positions outside cancer-type-enriched ATAC-seq peaks were enriched for pathways related to infection response, autophagy, and hormone signaling (**Fig. 3B and Table S6**), suggesting that nucleosome repositioning also occurs in regulatory contexts that are not readily detected by accessibility peaks.

**Fig. 3.**
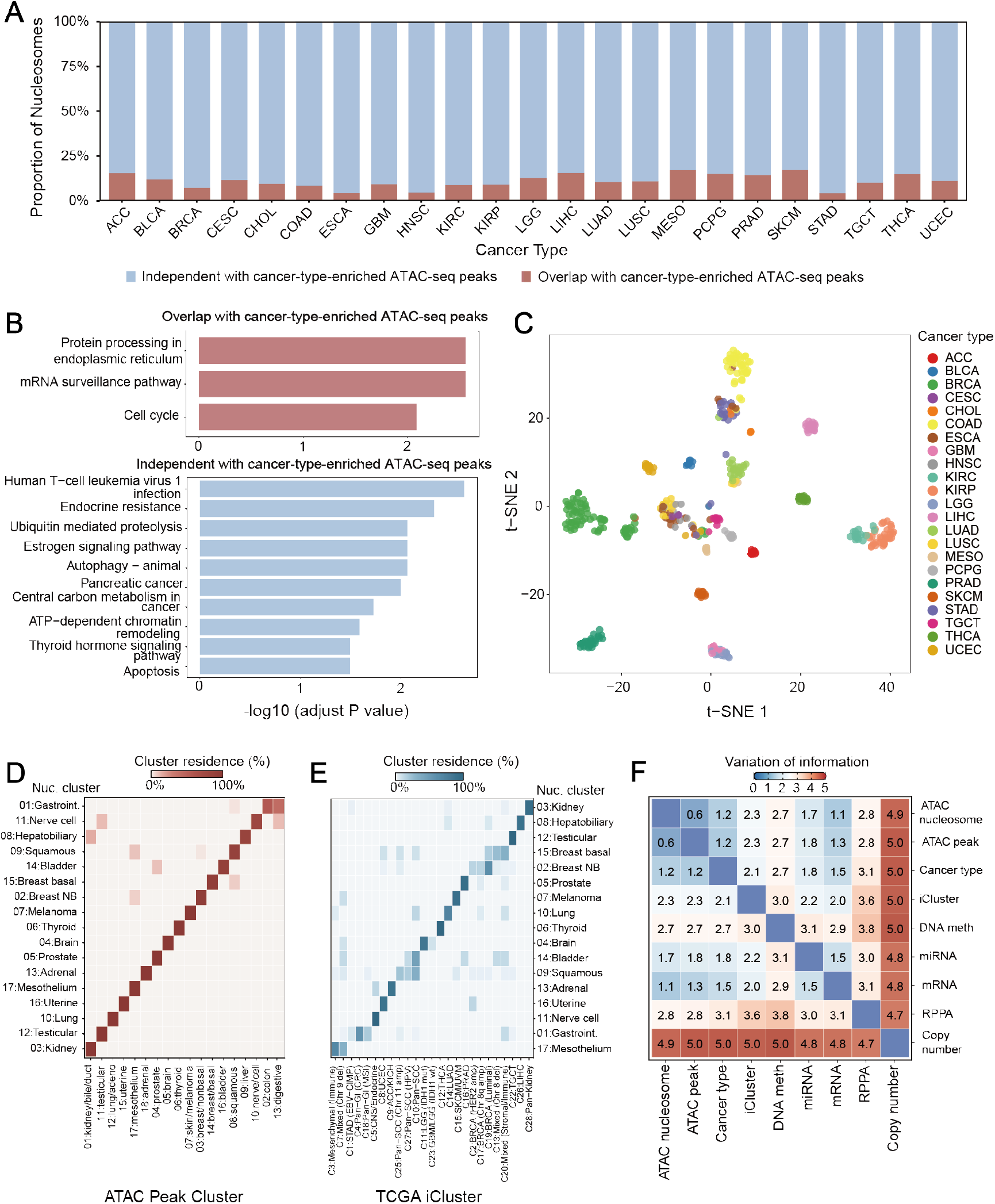
Cancer-type-enriched nucleosomes provide complementary information to chromatin accessibility for tumor stratification. **(A)** Proportion of cancer-type-enriched nucleosomes overlapping cancer-type-enriched ATAC-seq peaks across cancer types. Red and blue indicate nucleosomes overlapping and not overlapping ATAC-seq peaks, respectively. **(B)** Pathway enrichment analysis of genes associated with the top 1,000 cancer-type-enriched nucleosomes overlapping or not overlapping cancer-type-enriched ATAC-seq peaks, ranked separately within each group by mean nucleosome occupancy across all samples. **(C)** t-SNE visualization of 410 tumor samples from 23 cancer types based on cancer-type-enriched nucleosome occupancy profiles. Samples are colored by cancer type. **(D–E)** Residence heatmaps comparing nucleosome-based clustering with ATAC-seq peak-based clustering **(D)** and iCluster-based molecular subtypes **(E)**. **(F)** Heatmap showing the variation of information (VI) between nucleosome-based clustering and clustering based on other molecular features. Lower VI values indicate greater concordance between clustering results.

To determine whether these complementary nucleosome features capture biologically meaningful relationships among tumors, we performed unsupervised clustering based on the 250,000 cancer-type-enriched nucleosomes (**Fig. S5**), followed by t-distributed stochastic neighbor embedding (t-SNE) for visualization^39,40^. The resulting clusters not only separated individual cancer types but also preserved biologically meaningful relationships among tumors. For example, kidney renal papillary cell carcinoma (KIRP) clustered closely with kidney renal clear cell carcinoma (KIRC), consistent with their shared tissue origin, whereas lung squamous cell carcinoma (LUSC), esophageal carcinoma (ESCA), and head and neck squamous cell carcinoma (HNSC) occupied neighboring clusters, reflecting their common squamous histology (**Fig. 3C and Table S7**). These findings suggest that nucleosome positioning faithfully captures both cancer-type identity and broader biological relationships across cancers.

We next compared nucleosome-based clustering with previously reported molecular classifications, including ATAC-seq peak-based clustering^30^ and the multi-omics iCluster framework^41^. Nucleosome-based clustering showed substantial concordance with ATAC-seq peak-based clustering (**Fig. 3D**), consistent with their shared basis in chromatin organization. Importantly, however, nucleosome positioning resolved additional structure within 7 of 18 ATAC-defined clusters, affecting 231 samples (**Fig. S6**), and reorganized several biologically related cancer types. For example, KIRP, KIRC, and CHOL were grouped within a single ATAC-seq peak-defined cluster, whereas nucleosome-based clustering separated CHOL and placed it closer to LIHC. Similarly, STAD and COAD, which were separated in the peak-based analysis, clustered together based on nucleosome organization (**Fig. S6**). These changes indicate that nucleosome-based and accessibility-based clustering, despite their overall concordance, resolve distinct relationships among specific cancer types. In contrast, concordance with iCluster was comparatively lower (**Fig. 3E**), consistent with its integration of multiple molecular modalities extending beyond chromatin organization. Quantitative comparison using Variation of Information (VI), where lower values indicate greater similarity, further supported these relationships: nucleosome-based clustering was most similar to ATAC-seq peak-based clustering (VI = 0.6), followed by cancer-type classification (VI = 1.2), whereas substantially larger distances were observed relative to iCluster (VI = 2.3) and other molecular modalities (**Fig. 3F**). Together, these findings demonstrate that cancer-type-enriched nucleosome positioning preserves the core chromatin regulatory states captured by accessibility profiling while providing complementary structural information that refines tumor stratification.

### Nucleosome organization delineates molecular subtypes within tumors

Having established cancer-type nucleosome landscapes across cancers, we next investigated whether nucleosome positioning could resolve heterogeneity within individual tumor types. We first focused on breast invasive carcinoma (BRCA; n = 74), the largest cohort in our dataset. K-means clustering of variable nucleosomes’ occupancy identified three distinct subgroups, with the optimal cluster number determined by gap statistics (**Fig. 4A, Fig. S7A, and Table S8**). These nucleosome-defined subgroups exhibited heterogeneous distributions across pathological stage, estrogen receptor (ER) status, and progesterone receptor (PR) status (**Fig. 4B**), indicating that nucleosome profiles capture molecular heterogeneity not fully explained by these clinicopathological features. Notably, Subgroup 1 showed substantial overlap with the PAM50 Basal-like subtype (Fisher’s exact test, *P* = 2.031 × 10⁻¹³), an aggressive molecular subtype typically characterized by triple-negative breast cancer and poor clinical outcomes^42,43^ (**Fig. 4B**), suggesting that the Basal-like tumors exhibit a distinct nucleosome organization.

**Fig. 4.**
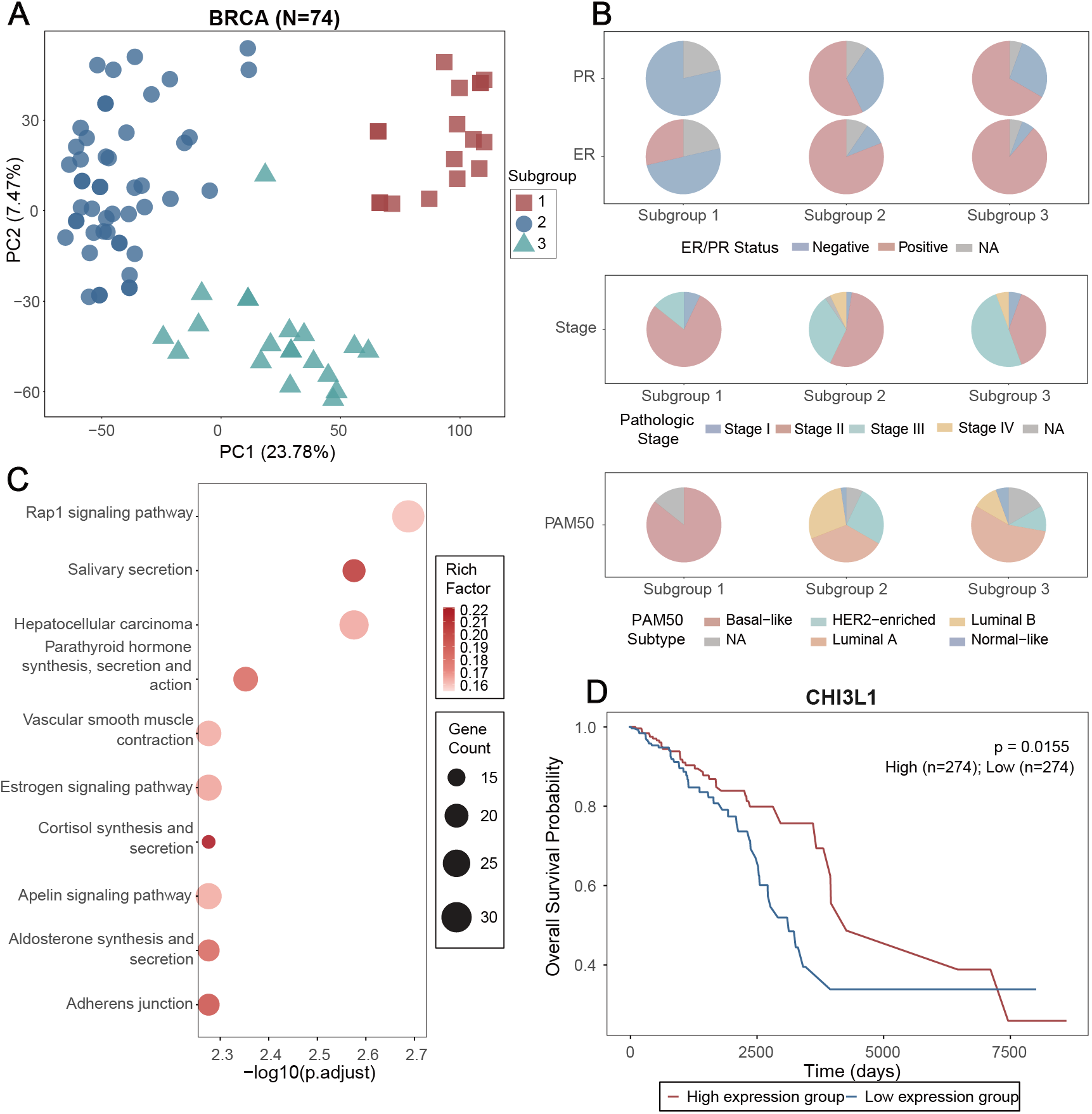
Nucleosome occupancy patterns reveal molecular heterogeneity within cancer types. **(A)** Principal component analysis (PCA) of BRCA samples based on nucleosome occupancy profiles. Each point represents an individual tumor sample and is colored according to its nucleosome-defined subgroup. **(B)** Pie charts showing the distributions of ER/PR status (top), pathlogical stages (middle), and PAM50 molecular subtypes (bottom) within each nucleosome-defined subgroup. **(C)** Functional enrichment analysis of genes associated with Subgroup 1-enriched nucleosomes in BRCA. **(D)** Kaplan–Meier analysis of overall survival in the TCGA BRCA stratified by CHI3L1 expression. Patients in the upper and lower expression quartiles were defined as the high- and low-expression groups, respectively; patients in the middle 50% were excluded from the analysis.

To characterize the regulatory programs underlying these subgroups, we performed functional enrichment analysis of genes associated with subgroup-enriched nucleosomes (**Table S9**). Subgroup 1 was significantly enriched for pathways linked to aggressive tumor behavior, including the Rap1 and estrogen signaling pathways^44–46^ (**Fig. 4C**), whereas the remaining subgroups were enriched for pathways characteristic of less invasive phenotypes (**Fig. S7B-C**). Among genes associated with promoter-localized subgroup-enriched nucleosomes, CHI3L1 contained the highest number of subgroup-enriched nucleosomes within its promoter region (**Fig. S7D**). Elevated CHI3L1 expression was significantly associated with poorer overall survival in the TCGA BRCA cohort (**Fig. 4D**), highlighting its potential clinical relevance in nucleosome-defined BRCA subgroups.

To assess the generalizability of this approach, we next analyzed kidney renal papillary cell carcinoma (KIRP; n = 34). Despite the smaller cohort size, clustering of variable nucleosomes similarly identified multiple molecular subgroups with distinct molecular characteristics (**Fig. S8A-B and Table S10**). We further identified SLC28A1 as a subgroup-enriched nucleosome-associated gene (**Table S11**), whose expression was significantly associated with patient survival (**Fig. S8C**). Although SLC28A1 has been implicated in cancer progression^47–49^, its potential role in KIRP has not, to our knowledge, been reported. Together, these findings demonstrate that nucleosome organization provides a robust framework for identifying clinically relevant tumor subtypes, uncovering subtype-specific regulatory programs, and nominating candidate prognostic biomarkers.

### The pan-cancer nucleosome atlas reveals cancer-associated nucleosome remodeling

We next investigated whether the pan-cancer nucleosome atlas could identify cancer-associated changes in nucleosome organization by comparing liver hepatocellular carcinoma (LIHC) with an independent normal liver reference cohort^50^. LIHC was selected for this analysis because of the availability of a large normal liver cohort (n = 187), enabling a well-powered tumor–normal comparison. After correcting for batch effects using RUVg^51^ (**Fig. S9**), we identified widespread differences in nucleosome occupancy between LIHC and normal liver tissues (**Table S12**). Differential nucleosomes were preferentially enriched at promoter regions (**Fig. 5A**; Chi-square test, *P* < 2.2e-16), indicating extensive reorganization of regulatory chromatin during hepatocarcinogenesis.

**Fig. 5.**
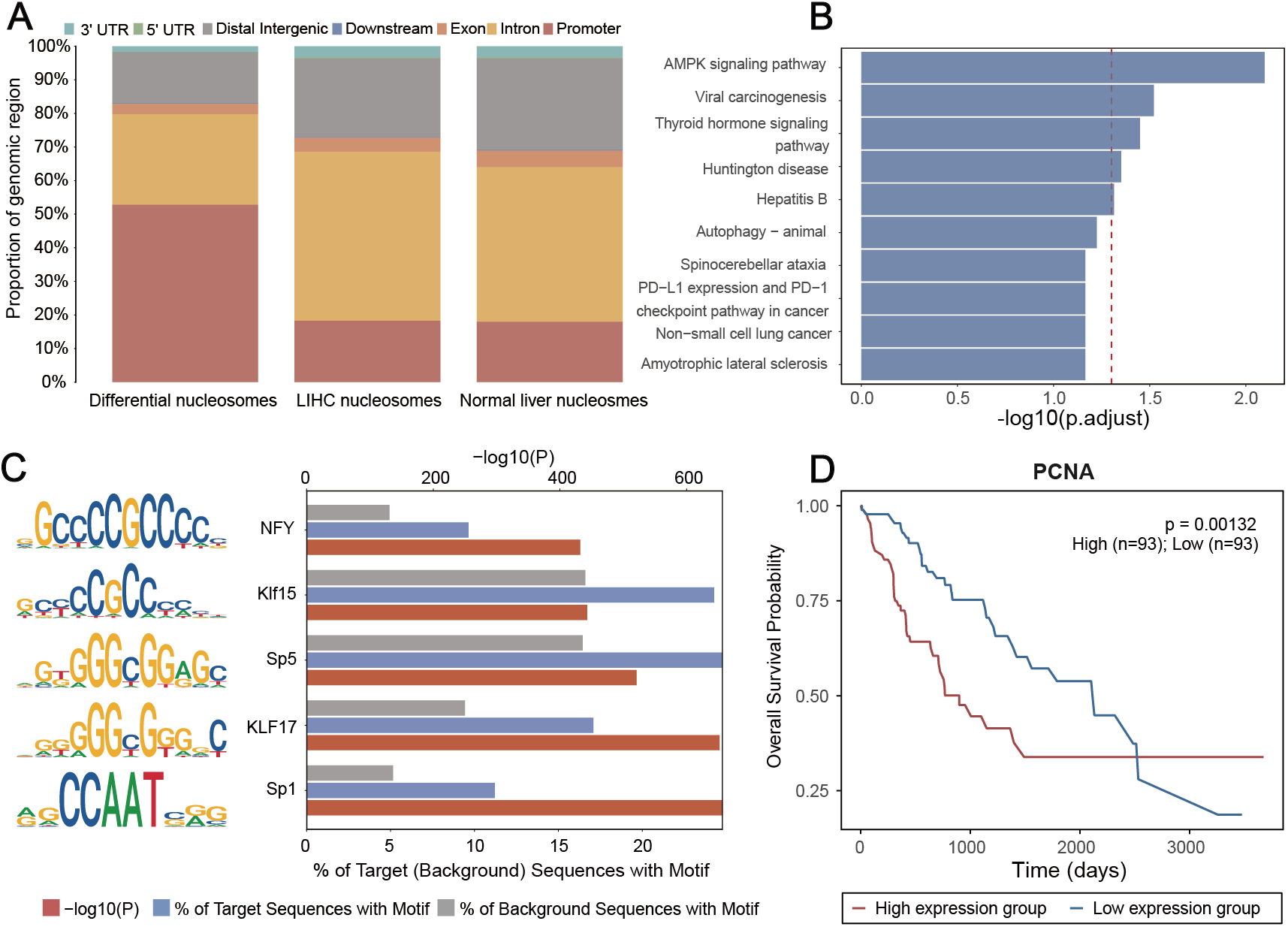
Cancer-associated nucleosome remodeling in hepatocellular carcinoma reveals regulatory and prognostic features. **(A)** Genomic distribution of nucleosomes identified in normal liver and LIHC tissues, and of differential nucleosomes identified between LIHC and normal liver samples. **(B)** Pathway enrichment analysis of genes associated with the top 1,000 differential nucleosomes, ranked by significance. The x-axis indicates −log10(adjusted P value). **(C)** Transcription factor (TF) motif enrichment analysis of differential nucleosome regions. The left panel shows sequence logos for the five most significantly enriched TF motifs, and the right panel shows motif enrichment significance [−log10(P value)] together with the frequencies of each motif in differential nucleosome and background regions. **(D)** Kaplan–Meier analysis of overall survival in the TCGA LIHC cohort stratified by PCNA expression. Patients in the upper and lower expression quartiles were defined as the high- and low-expression groups, respectively; patients in the middle 50% were excluded from the analysis.

Genes associated with differential nucleosomes were enriched in pathways implicated in liver cancer, including AMPK signaling and autophagy (**Fig. 5B**), both of which play key roles in metabolic reprogramming and tumor progression in LIHC^52,53^. Transcription factor motif analysis further revealed significant overrepresentation of binding sites for SP1, NF-Y, and members of the KLF family (**Fig. 5C and Table S13**), key regulators of cell proliferation, invasion, and tumor progression^54,55,56^, suggesting that nucleosome remodeling is associated with oncogenic regulatory programs in LIHC. At the gene level, two of the most significant promoter-associated differential nucleosomes mapped to PCNA and ARL4D. PCNA is a key regulator of DNA replication and repair^57^ and elevated PCNA expression was significantly associated with poorer overall survival in the TCGA LIHC cohort (**Fig. 5D**). In contrast, ARL4D has not been previously implicated in LIHC, highlighting it as a potential candidate for further investigation.

Collectively, these findings demonstrate that the pan-cancer nucleosome atlas not only enables systematic identification of cancer-associated nucleosome remodeling but also reveals the regulatory pathways and transcriptional networks. These results highlight the value of nucleosome positioning as a complementary layer of epigenetic information for dissecting the regulatory mechanisms of human cancers.

### Tissue-derived nucleosome signatures are preserved in cfDNA and inform cancer classification and detection

Given that cfDNA retains nucleosome organization from its tissue of origin^15–18^, we next investigated whether cancer-type-enriched nucleosome signatures could distinguish tumor origins from cfDNA. Using a cfDNA whole-genome sequencing dataset comprising patients with BRCA, colon adenocarcinoma (COAD), and stomach adenocarcinoma (STAD)^16^, we quantified cfDNA occupancy at cancer-type-enriched nucleosomes and trained a multi-class classifier. Ten-fold cross-validation yielded a micro-averaged area under the receiver operating characteristic curve (AUROC) of 0.877 (95% CI: 0.838–0.918) for distinguishing each cancer type from the others (**Fig. 6A**), with overall and balanced accuracies of approximately 0.75 (**Fig. S10**). Genes associated with the selected nucleosome features were enriched in cancer-relevant pathways (**Fig. 6B and Table S14**), including cell polarity pathways in BRCA^58^, nucleotide biosynthesis in COAD^59^ and maintenance of gastrointestinal epithelium in STAD^60^, supporting the biological relevance of these cfDNA nucleosome signatures.

**Fig. 6.**
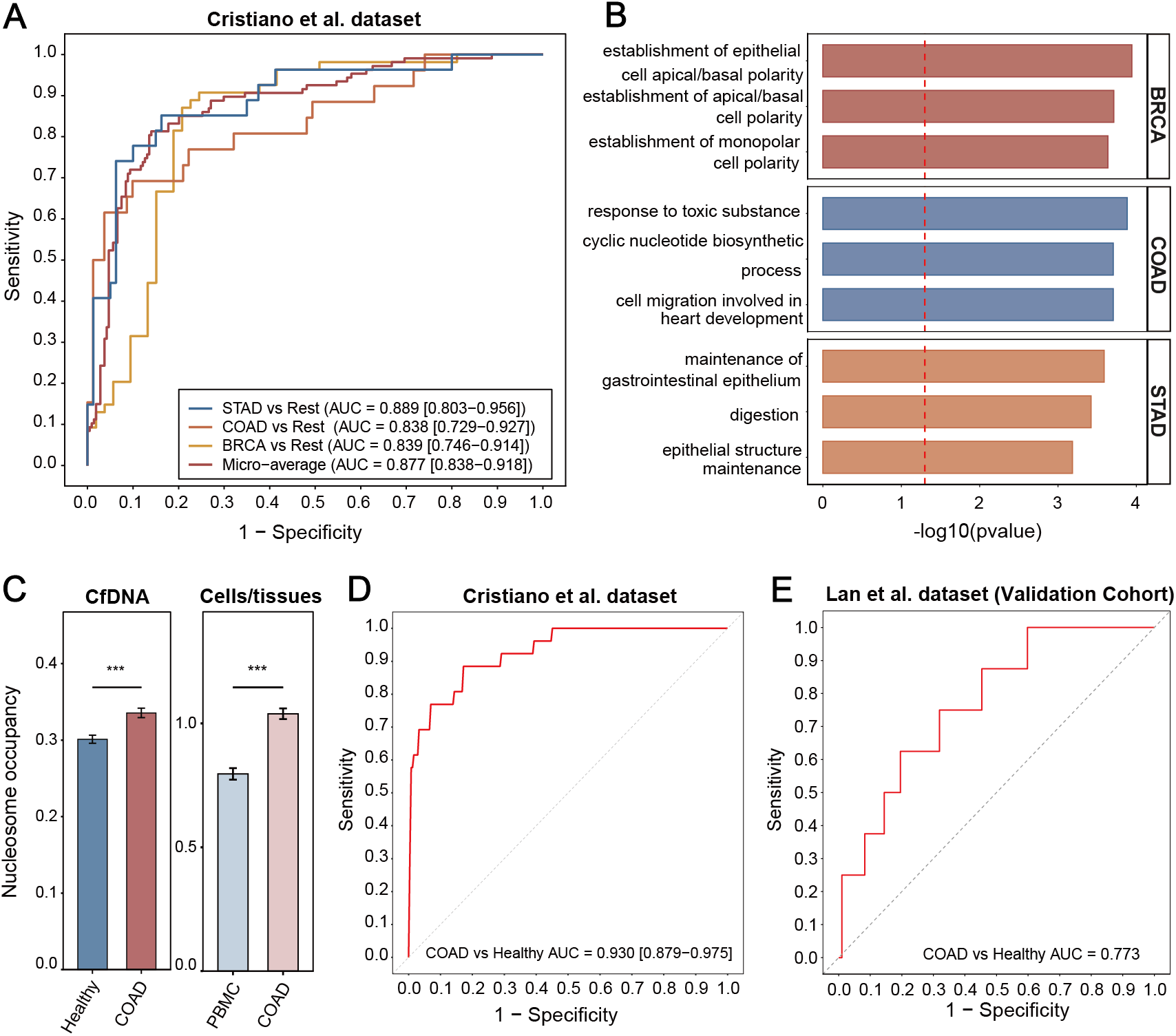
Tissue-derived nucleosome signatures enable cfDNA-based cancer classification and detection. **(A)** Multiclass classification of COAD (n = 26), BRCA (n = 54), and STAD (n = 27) using cfDNA occupancy at cancer-type-enriched nucleosomes derived from tumor ATAC-seq data. Performance is shown by class-specific and micro-averaged receiver operating characteristic (ROC) curves. The 95% confidence interval (CI) was estimated using 2,000 bootstrap replicates and shown in the parentheses. **(B)** Pathway enrichment analysis of genes associated with the nucleosome features selected for BRCA, STAD, and COAD in the multiclass classification analysis. **(C)** Barplot showing the concordance of tissue-derived differential nucleosome signals with cfDNA nucleosome occupancy in COAD. For each differential nucleosome, occupancy was averaged across samples within each group. Bars represent the mean occupancy across differential nucleosomes, and error bars indicate the standard error of the mean (SEM). **(D)** ROC analysis for cfDNA-based detection of COAD using tissue-derived differential nucleosome features. Model performance was evaluated by stratified 10-fold cross-validation in the cohort comprising 26 patients with COAD and 244 healthy individuals (Cristiano et al. dataset from FinaleDB). The 95% confidence interval (CI) was estimated using 2,000 bootstrap replicates and shown in the parentheses. **(E)** ROC analysis for independent validation of the COAD classifier in the HRA004005 cfDNA cohort comprising 8 patients with COAD and 97 healthy individuals. The SVM classifier trained on the discovery cohort (Cristiano et al. dataset from FinaleDB) was applied to the independent validation cohort without model retraining.

We next investigated whether tumor-associated differential nucleosomes identified in tissue could also be detected in circulation. We focused on COAD, which had a relatively large sample size in our atlas (n = 38) and, importantly, an independent cfDNA cohort was available for validation. By comparing tumor nucleosome profiles with those of peripheral blood mononuclear cells (PBMC), the major source of cfDNA in healthy individuals^61^, we identified 16,391 nucleosomes with significantly altered occupancy in COAD (**Fig. S11 and Table S15**). These tissue-derived alterations showed concordant occupancy changes in an independent cfDNA cohort comprising 244 healthy controls and 26 COAD patients^16^ (**Fig. 6C**), indicating that tumor-associated nucleosome remodeling is detectable in circulating DNA.

We then evaluated the utility of these tissue-derived differential nucleosomes for cancer detection. The COAD classifier achieved an AUROC of 0.930 (95% CI: 0.879–0.975) under ten-fold cross-validation (**Fig. 6D**) and an AUROC of 0.773 in an independent validation cohort^62^ (n = 105; **Fig. 6E**), supporting the generalizability of tissue-derived nucleosome signatures across cohorts. Nucleosome features retained in the final classifiers localized near cancer-relevant genes (**Table S16**), including PMEPA1^63^, KLF5^64^, and JAM2^65^. These findings indicate that tissue-derived nucleosome alterations are detectable in cfDNA and can provide biologically informed features for noninvasive cancer detection.

Collectively, these results demonstrate that nucleosome signatures derived from the pan-cancer atlas are preserved in circulating cfDNA and can be leveraged for both cancer classification and detection. More broadly, they establish the pan-cancer nucleosome atlas as a translational resource linking tissue-specific chromatin organization with noninvasive liquid biopsy biomarkers.

### OncoNucAtlas enables interactive exploration of pan-cancer nucleosome landscapes

To facilitate access to and reuse of the pan-cancer nucleosome atlas, we developed OncoNucAtlas, an interactive web resource integrating sample-level, cancer-type-level, and cancer-type-enriched nucleosome maps across 410 samples from 23 cancer types (**Fig. S12**). OncoNucAtlas enables users to query annotated nucleosome positions and occupancy profiles, compare nucleosome occupancy across cancer types, download sample-level nucleosome maps, and visualize nucleosome occupancy tracks through the UCSC Genome Browser. Together, these functions provide an accessible platform for exploring and reusing nucleosome organization across human cancers.

## Discussion

Chromatin accessibility has become a central framework for studying cancer epigenomes^30^, yet the accompanying organization of nucleosomes across human cancers has remained incompletely characterized. Here, we constructed the first pan-cancer nucleosome atlas spanning 23 tumor types using high-quality TCGA ATAC-seq data and demonstrated that nucleosome positioning can be robustly inferred from nucleosome-sized ATAC-seq fragments. Validation with independent MNase-seq datasets confirmed the accuracy of the inferred nucleosome occupancy patterns, while the well-phased organization around transcription start sites and CTCF binding sites further supports the biological fidelity of the atlas. Together, these results establish a comprehensive reference of nucleosome organization across human cancers and provide a resource for investigating chromatin architecture at both shared and cancer-specific regulatory elements.

A major finding of this study is that nucleosome positioning provides regulatory information beyond conventional chromatin accessibility analyses. Although ATAC-seq peaks have become the standard representation of accessible chromatin, more than two-thirds of cancer-type-enriched nucleosome positions identified in our study were not captured by ATAC-seq peaks, indicating that changes in nucleosome organization reveal regulatory events that are largely invisible to peak-based analyses. Moreover, cancer-type-enriched nucleosomes were associated with lineage-specific transcription factor networks and biological pathways, whereas conserved nucleosome positions preferentially localized to promoters and genes involved in core cellular functions. Together, these observations suggest that nucleosome positioning captures both the dynamic chromatin remodeling underlying tumor identity and the conserved regulatory architecture required for malignant cell survival.

Importantly, we demonstrate the translational potential of tissue-derived nucleosome positioning by showing that atlas-derived nucleosome signatures are preserved in circulating cell-free DNA. Cancer-type-enriched nucleosome landscapes enabled accurate classification of tumor origin, whereas tissue-derived differential nucleosome signatures supported cancer detection in independent cfDNA cohorts. These results reinforce accumulating evidence that cfDNA fragmentation patterns retain the chromatin organization of their cells of origin and further establish nucleosome positioning as a biologically interpretable structural biomarker for liquid biopsy. As sequencing costs continue to decline and cfDNA profiling becomes increasingly integrated into clinical workflows^66,67^, atlas-guided nucleosome signatures may provide a complementary strategy for noninvasive cancer diagnosis and tissue-of-origin prediction.

Several limitations should be acknowledged. First, because nucleosome positions were inferred from ATAC-seq rather than measured directly, the resulting atlas is inherently biased toward accessible chromatin regions where Tn5 transposase efficiently inserts. Consequently, nucleosome organization within constitutively compact heterochromatin is likely underrepresented compared with approaches such as MNase-seq^22^ or chemical cleavage-based mapping^68^. Nevertheless, these accessible regions encompass the majority of functional regulatory elements^69,70^ and therefore remain highly informative for investigating cancer-associated chromatin remodeling. Second, sample sizes varied considerably across cancer types, particularly for rare tumors, which may influence the robustness of cancer-type-enriched nucleosome identification despite the normalization and quality-control procedures applied in this study. Continued expansion of cancer epigenomic datasets will further improve the completeness and statistical power of the atlas.

In conclusion, our study establishes a pan-cancer nucleosome atlas that complements existing chromatin accessibility resources by providing a genome-wide view of nucleosome organization across human cancers. Beyond serving as a reference map, the atlas reveals an additional layer of epigenetic regulation that contributes to cancer-specific transcriptional programs, tumor heterogeneity, and conserved oncogenic processes, while enabling the development of nucleosome-based liquid biopsy biomarkers. Future integration with higher-resolution nucleosome mapping technologies^71^, single-cell chromatin profiling^72^, and multi-omics datasets will further elucidate the role of nucleosome organization in tumor evolution and accelerate its translation into precision oncology.

## Methods

### Data collection

ATAC-seq data from 410 samples across 23 cancer types were downloaded from The Cancer Genome Atlas (TCGA) through the Genomic Data Commons (https://portal.gdc.cancer.gov/) portal. To validate nucleosome profiles inferred from ATAC-seq, four publicly available datasets containing matched ATAC-seq and MNase-seq data were downloaded from the Gene Expression Omnibus (GEO) database: GSE124903 (TTC1240, malignant rhabdoid tumor cell line), GSE160470 (Kasumi-1, acute myeloid leukemia cell line), GSE201801 (MDA-MB-231, triple-negative breast cancer cell line), and GSE144454 (eHAP, near-haploid human cell line). In addition, MNase-seq data from COAD and LUAD were collected from GSE74340 for validation of tissue-derived nucleosome profile. Normal liver ATAC-seq data were obtained from GSE277774, with two samples exhibiting abnormal fragment-size distributions excluded from downstream analyses. ATAC-seq data of PBMCs were obtained from GSE193140. Whole-genome sequencing (WGS) data of cfDNA from Cristiano *et al.* (samples EE87811 to EE88182) and Jiang *et al.* (samples EE85756 to EE86002) were downloaded from FinaleDB^73^, whereas the HRA004005 dataset downloaded from the Genome Sequence Archive for Human (GSA-Human) was used for independent validation. Detailed sample information for all datasets is summarized in **Table S1**.

### Raw ATAC-seq and MNase-seq data processing

For all ATAC-seq and MNase-seq data, raw FASTQ files were first processed with Trim Galore to remove adapter sequences and low-quality reads^74^ (version 0.6.10; -- paired --fastqc). The resulting clean reads were aligned to the human reference genome (hg38) using Bowtie2^75^ (version 2.5.4, --very-sensitive -X 2000). Aligned reads were sorted and filtered using SAMtools^76^, retaining only properly paired, uniquely mapped reads with a mapping quality score greater than 30 while removing supplementary alignments (version 1.18; -f 2 -q 30 -F 2048). PCR duplicates were subsequently removed using Picard MarkDuplicates^77^.

### Construction of the pan-cancer nucleosome atlas

In this study, we focused on two widely-used features of the nucleosome landscape: nucleosome positioning, which refers to precise genomic locations of nucleosomes, and nucleosome occupancy, which reflects the relative abundance of nucleosomes at a given genomic region.

- Identification of sample- and cancer-type-level nucleosome sets For each ATAC-seq dataset, nucleosome positions were identified using deNOPA^25^ and NucleoATAC^24^ with default parameters, respectively. DeNOPA was run with the parameter *“-c chr[1-9][0-9]{,1}|chrX*” to retain nucleosome calls from autosomes and chromosome X. For NucleoATAC, broad open chromatin regions were first identified using MACS2^78^ (version 2.2.9.1; *--shift -75 --extsize 150 --nomodel --nolambda --keep-dup all --broad --broad-cutoff 0.01 -g 2747877702*). The resulting broad peaks were extended by 1 kb in both directions and merged into non-overlapping regions using BEDTools^79^. NucleoATAC was then run on these merged regions using default parameters. For each method, the reported nucleosome dyads were extended by 73 bp upstream and downstream to generate 147-bp nucleosome intervals corresponding to canonical nucleosome footprints. Intervals overlapping ENCODE blacklist regions^80^ or extending beyond chromosome boundaries were subsequently excluded. Nucleosome occupancy was quantified by extracting ATAC-seq fragments between 101 to 215 bp in length, representing mono-nucleosome-protected fragments. Fragments whose centers fell within ±50 bp of each nucleosome interval were used to calculate occupancy profiles using DANPOS3^81^ (*-m 1 -q 1 -jd 73 -n N*). To account for the differences in sequencing depth and library size across samples, raw occupancy profiles were normalized by dividing by the total number of mapped reads in the original BAM file and multiplied by 1,000,000 to generate the normalized occupancy score. Because extending nucleosome dyads to 147-bp intervals can generate overlapping nucleosome regions, an greedy overlap-resolution procedure was applied to each sample independently. Nucleosomes were first ranked in descending order according to their occupancy scores. The highest-scoring nucleosome was retained, and any nucleosome that directly overlapped with this nucleosome was removed. Then, this procedure was repeated iteratively with the next highest-scoring nucleosome until all nucleosomes had either been retained or removed due to overlap with a higher-scoring nucleosome. By resolving only direct overlaps at each iteration, this greedy procedure avoids the unnecessary removal of nucleosomes that do not directly overlap but are connected through intermediate overlapping intervals, thereby generating a set of fixed-width, non-overlapping nucleosome intervals. The resulting collection of nucleosomes for each sample was defined as the sample-level nucleosome set. To construct a cancer-type-level consensus nucleosome set, sample-level nucleosomes within each cancer type were pooled separately for deNOPA and NucleoATAC. Overlapping nucleosomes across samples were resolved using the greedy overlap-resolution procedure described above, generating a nonredundant candidate set for each cancer type and caller. Each sample-level nucleosome set was then intersected with the corresponding candidate set, and a candidate nucleosome was considered supported by a sample if their 147-bp intervals overlapped. Candidate nucleosomes supported by at least two independent samples were retained as reproducible cancer-type-level nucleosomes, reducing the influence of sample-specific artifacts and low-confidence nucleosome calls. Finally, the cancer-type-level nucleosome sets independently identified by deNOPA25 and NucleoATAC were intersected, requiring an overlap of ≥73 bp. For each overlapping pair, the midpoint of the overlapping region was defined as the consensus dyad and extended by 73 bp on either side to generate a 147-bp interval. These intervals constituted the final high-confidence cancer-type-level nucleosome set for each cancer type (**Fig. S3**).
- Identification of cancer-type-enriched and shared nucleosomes across cancer types To identify cancer-type-enriched nucleosomes, mean nucleosome occupancy across samples was first calculated for each cancer-type-level nucleosome and normalized using the “score per million” strategy^30^. For each cancer type, the occupancy score of each nucleosome was divided by the sum of occupancy scores across all nucleosomes and multiplied by 10⁶, thereby facilitating comparison of nucleosome occupancy across cancer types. The normalized cancer-type-level nucleosome sets from all 23 cancer types were then pooled while retaining their cancer-type labels. Overlapping nucleosomes were resolved using the greedy procedure described above, with the nucleosome exhibiting the highest normalized occupancy retained together with its corresponding cancer-type label. The resulting nonredundant collection of nucleosomes was defined as the cancer-type-enriched nucleosome set (**Fig. S3**). To identify nucleosomes shared across cancer types, the final high-confidence cancer-type-level nucleosome sets were intersected across all 23 cancer types, and only nucleosome intervals shared by all cancer types were retained. The midpoint of each shared region was defined as the consensus dyad and extended by 73 bp on either side, generating a 147-bp interval. These nucleosomes were designated as pan-cancer nucleosomes (**Fig. S3**).

### Nucleosome positioning and occupancy analysis for MNase-seq

For MNase-seq data, nucleosome positioning was determined using the *dpos* module of DANPOS3 (*-m 1 -n F*), and the resulting nucleosome dyad positions were defined as nucleosome positioning. Nucleosome occupancy was quantified using the *dpos* module of DANPOS3 (*-m 1 -q 1 -jd 73 -c 10000000*). The resulting normalized occupancy scores were defined as nucleosome occupancy.

### Motif enrichment analysis

Motif enrichment analysis was performed using HOMER^82^ (*findMotifsGenome.pl*, - *mset vertebrates*), with 147-bp nucleosome intervals as input. Motif enrichment was evaluated by comparing motif frequencies between target and background regions using HOMER’s default statistical framework. For **Fig. 2E**, motif enrichment was analyzed separately for pan-cancer nucleosomes and cancer-type-enriched nucleosomes from each cancer type. A common background comprising the union of cancer-type-enriched nucleosome regions across all 23 cancer types was used to facilitate comparison across analyses. For pan-cancer nucleosomes, the 20 most significantly enriched motifs ranked by hypergeometric test P values were visualized. For cancer-type-enriched nucleosomes, we focused on 15 cancer-type-associated transcription factor motifs previously highlighted by Corces *et al.*^30^ with corresponding entries in the HOMER motif database for visualization. For **Fig. 5C**, differential nucleosome regions between normal and LIHC tissues were used as target regions, with non-differential nucleosomes as background. Motif enrichment was performed using the same HOMER parameters, and the five most significantly enriched motifs ranked by hypergeometric test P values were visualized.

### Nucleosome-based clustering and visualization

Cancer-type-enriched nucleosome intervals were used for clustering analysis. For each sample, normalized nucleosome occupancy values were log2-transformed to reduce the influence of highly abundant signals and subsequently quantile-normalized across samples using the *normalizeBetweenArrays* function from the *limma* package^83^. Nucleosome intervals were ranked by their variance across samples using *matrixStats* package^84^. Principal component analysis (PCA) was performed using the *prcomp_irlba* function from the *irlba* package^85^ with 50 PCs initially computed (center = TRUE, scale. = FALSE). Density-based clustering was performed on the PCA representation using *DensityClust*^40^, with *rho* and *delta* parameters used to define cluster centers.

Clustering parameters were selected by systematically evaluating the number of variable nucleosome intervals, the number of retained PCs, and the *DensityClust* parameters. The *rho* parameter was first evaluated using *DensityClust* decision plots based on the 250,000 most variable nucleosomes and principal components explaining 85% of the cumulative variance, and *rho = 2* was selected based on the separation of candidate cluster centers (**Fig. S5A**). With *rho* fixed at 2 and 250,000 nucleosomes as input, cumulative explained-variance thresholds of 80%, 85%, 90%, and 95% were evaluated, together with *delta* values ranging from 100 to 150. The parameter combination yielding the highest silhouette coefficient was selected (*delta* = 105; cumulative explained-variance = 85%) (**Fig. S5B**). Finally, the number of input nucleosomes was subsequently evaluated from 100,000 to 500,000 in increments of 50,000 while holding the other parameters fixed, with the 250,000 showing the highest silhouette coefficient (**Fig. S5C**). The final clustering therefore used the 250,000 most variable nucleosome intervals, PCs explaining 85% of the cumulative variance, *rho* = 2, and *delta* = 105.

For visualization, t-distributed stochastic neighbor embedding (t-SNE) was performed on the 250,000 most variable nucleosome intervals using the Rtsne package^86^ (v0.17), with a perplexity of 20 and 10,000 iterations. t-SNE was applied to the normalized occupancy profiles of the 250,000 most variable nucleosomes and was used solely for visualization; samples were colored according to cluster assignments obtained independently from the PCA-based *DensityClust* analysis.

### Cancer subtyping analysis

Nucleosome occupancy at cancer-type-level nucleosome positions was quantified for each sample from the corresponding BAM files using the strategy described above. Briefly, occupancy was calculated from nucleosome-sized fragments, normalized by the total number of mapped reads per sample, log2-transformed, and quantile-normalized across samples.

For each cancer type, unsupervised clustering was performed using the normalized nucleosome occupancy matrix. The 25,000 nucleosomes with the highest variance across samples were selected, and their occupancy values were standardized as row-wise z-scores. The optimal number of clusters (k) was determined using the gap statistic with the *firstSEmax* criterion implemented in the *cluster::maxSE*. K-means clustering was then performed using the selected *k,* with 25 random initializations (*nstart = 25*). The resulting clusters were designated as nucleosome-defined tumor subgroups.

To identify subgroup-enriched nucleosomes, a one-versus-rest strategy was applied. For each subgroup, nucleosome occupancy was compared between the target subgroup and all remaining subgroups using a two-sided Wilcoxon rank-sum test, followed by Benjamini-Hochberg correction for multiple testing. Nucleosomes with a false discovery rate (FDR) ≤ 0.05 and absolute log2 fold change ≥ 1 were defined as subgroup-enriched nucleosomes. These nucleosomes were annotated to their nearest genes using ChIPseeker^87^, and the associated genes were subjected to functional enrichment analysis to characterize biological pathways distinguishing each subgroup.

To assess the clinical relevance of genes associated with subgroup-enriched nucleosomes, RNA-seq expression and overall survival (OS) data were obtained from TCGA using TCGAbiolinks^88^ and UCSCXenaTools^89^ packages, respectively. Representative genes were prioritized based on prominent promoter-associated subgroup-enriched nucleosome signals, including promoter nucleosome density and the magnitude of occupancy differences between subgroups. For each candidate gene, patients in the upper and lower quartiles of gene expression were classified as high- and low-expression groups, respectively, with the middle 50% excluded from survival analysis. Kaplan–Meier survival curves were generated using the survival and survminer packages^90^, and differences in OS between the two groups were evaluated using the log-rank test.

### Identification of differential nucleosomes between LIHC and normal liver tissues

ATAC-seq data from normal liver tissues (GSE277774) were processed using the same nucleosome-calling procedure described above. Briefly, sample-level nucleosomes were independently identified using deNOPA and NucleoATAC and consolidated across normal liver samples using the greedy overlap-resolution procedure. The resulting caller-specific sets were intersected, requiring an overlap of ≥73 bp, to generate the final high-confidence normal liver nucleosome set.

For tumor-normal comparison, the normal liver nucleosome set was merged with the LIHC nucleosome set to generate a union set of nucleosome positions. Nucleosome occupancy was quantified for each ATAC-seq sample using DANPOS3 based on nucleosome-sized fragments (101–215 bp), as described above. The resulting occupancy matrix was upper-quartile normalized and log2-transformed before batch-effect correction and differential analysis.

To identify stable negative-control nucleosomes for RUVg-based batch correction^51^, we leveraged independent cfDNA datasets. Because cfDNA contains nucleosome-derived signals originating from multiple tissues, we reasoned that nucleosome positions showing consistently stable occupancy across cancer and healthy cfDNA samples would provide a set of relatively invariant loci suitable as negative controls for estimating unwanted technical variation. This external strategy also avoided defining negative controls directly from the LIHC and normal liver ATAC-seq datasets used for differential analysis. cfDNA nucleosome occupancy was quantified at the above identified union nucleosome positions using DANPOS3 and cfDNA fragments (80–200 bp) (Jiang *et al.* dataset, healthy and LIHC samples EE85756 to EE86002 from FinaleDB), followed by “score per million” normalization described above. Nucleosomes were then filtered for reliable detection in both tissue and cfDNA datasets. For tissue ATAC-seq data, nucleosomes were required to have normalized occupancy greater than zero in at least 50% of both LIHC and normal liver samples. For cfDNA data, nucleosomes were required to have a sequencing depth of at least three reads in ≥80% of both cancer and healthy samples. Among the retained nucleosomes, the 25% showing the lowest coefficient of variation across cfDNA samples were selected as negative controls for RUVg.

Subsequently, RUVg was applied to the tissue ATAC-seq occupancy matrix using these negative-control nucleosomes to estimate factors representing unwanted variation. The inferred factors were included as covariates in the probabilistic index model (PIM)^91^ for differential nucleosome analysis between LIHC and normal liver tissues. P values were adjusted using the Benjamini–Hochberg procedure, and nucleosomes with FDR < 0.05 were considered differential.

### cfDNA-based cancer detection using differential nucleosomes between tumor and PBMCs

- Identification of differential nucleosomes between tumor and PBMCs To identify cancer-associated nucleosome features for cfDNA-based cancer detection, we focused on COAD, which had relatively large sample sizes in the pan-cancer ATAC-seq cohort. Differential nucleosomes were identified by comparing COAD tissues with PBMCs (GSE193140). A union nucleosome set was generated by combining the COAD cancer-type-level nucleosome set with the PBMC nucleosome set, which was constructed using the same procedure described above for normal liver tissues. Nucleosome occupancy was quantified at these union positions in each ATAC-seq sample using DANPOS3 and nucleosome-sized fragments. Raw occupancy values were upper-quartile normalized and log2-transformed as described above. Batch-effect correction and differential analysis were performed using the same framework applied to the LIHC–normal liver comparison. Briefly, cfDNA nucleosome occupancy for healthy and COAD samples was quantified at the union nucleosome positions using DANPOS3 and cfDNA fragments of 80–200 bp (Cristiano *et al.* dataset, samples EE87811 to EE88182 from FinaleDB). Nucleosomes showing stable occupancy across cfDNA samples (coefficient of variation < 0.5) were used as negative controls for RUVg to estimate unwanted variation. The resulting RUVg factors were incorporated as covariates into a PIM to identify differential nucleosomes between COAD and PBMCs. P values were adjusted using the Benjamini–Hochberg procedure, and nucleosomes with FDR < 0.05 were retained as candidate features for cfDNA-based cancer detection.
- Machine learning model construction To evaluate whether tumor-derived differential nucleosome features could distinguish patients with cancer from healthy individuals, nucleosome occupancy at the differential nucleosome positions was quantified in cfDNA samples (Cristiano *et al.* dataset, samples EE87811 to EE88182 from FinaleDB), and a linear support vector machine (SVM) implemented in the *e1071* package was used for binary classification^92^. Differential nucleosomes identified independently from the COAD–PBMC ATAC-seq comparison were first filtered based on sequencing coverage in the Cristiano *et al.* cfDNA cohort, retaining nucleosomes with a coverage of ≥3 reads in >80% of both healthy and COAD samples. Model performance was evaluated using stratified 10-fold cross-validation. Within each cross-validation iteration, feature selection and preprocessing were performed exclusively using the training samples to prevent information leakage. The retained differential nucleosomes were ranked according to the absolute Pearson correlation between their cfDNA occupancy values and cancer status within the training set, and the top 50 nucleosomes were selected. Feature values were used without any scaling or standardization. The SVM was trained on the resulting training matrix and used to predict cancer status in the corresponding test set. Prediction scores from held-out samples were used to construct receiver operating characteristic (ROC) curves and calculate the area under the ROC curve (AUROC) using the pROC package. The COAD classifier was further evaluated in an independent cfDNA WGS cohort (HRA004005). For this external validation, the final model was trained using the complete discovery cohort from Cristiano *et al.* dataset. The 50 differential nucleosome features with the highest absolute Pearson correlation with cancer status in the discovery cohort were selected. Nucleosome occupancy at the same feature positions was quantified in the independent validation samples using the same cfDNA processing and occupancy-quantification procedure described above. Feature values were directly entered into the pretrained COAD classifier without scaling, feature selection, or model fitting. Prediction scores were used to calculate the AUROC in the independent validation cohort using the pROC package.

### cfDNA-based cancer classification using cancer-type-enriched nucleosomes

To evaluate whether cancer-type-enriched nucleosome signatures could discriminate cancer types in cfDNA, we developed a multi-class classification framework for COAD, BRCA, and STAD. Among the four cancer types represented in both the TCGA-derived nucleosome reference and the cfDNA dataset (Cristiano *et al.* dataset from FinaleDB), these three had the largest sample sizes in both datasets and were therefore selected for analysis. Cancer-type-enriched nucleosomes identified from tumor tissue ATAC-seq data and assigned to each corresponding cancer type were used as candidate features.

A SVM implemented in the e1071 package was used for multiclass classification, and model performance was evaluated using stratified 10-fold cross-validation. Within each cross-validation fold, feature selection and preprocessing were performed exclusively using the training samples to prevent information leakage. For each cancer type, only nucleosomes previously identified as enriched for that cancer type were considered as candidate features. These features were ranked using a one-versus-rest strategy according to the absolute Pearson correlation between cfDNA nucleosome occupancy and the corresponding binary class label within the training set, and the top 30 features for each cancer type were selected. The resulting 90 cancer-type-enriched candidate features were combined into a single feature matrix and standardized using the mean and standard deviation estimated from the training samples; the same transformation was then applied to the corresponding held-out test samples.

The SVM model was trained using the resulting training data and used to predict cancer type in the held-out samples. Multiclass classification performance was evaluated using overall accuracy and balanced accuracy according for class imbalance. In addition, one-versus-rest ROC analysis was performed for each cancer type using out-of-fold prediction scores. Micro-averaged ROC analysis was performed by pooling the one-versus-rest predictions across classes.

### OncoNucAtlas database construction

OncoNucAtlas (https://liymlab.jysw.suda.edu.cn/OncoNucAtlas) was implemented as a web-based platform comprising a React frontend, a FastAPI backend, and a PostgreSQL database, and was deployed using Docker containers. All genomic coordinates were standardized to the hg38 reference genome. Processed nucleosome data were organized at three levels—sample-level, cancer-type-level, and cancer-type-enriched nucleosomes—and integrated with genomic, gene, and transcript annotations using the R package ChIPseeker (version 1.42.0). Sample-level nucleosome occupancy values were linked to the corresponding nucleosome records to enable comparison of occupancy patterns across individual samples and cancer types.

Database indexing enables efficient paginated queries by gene symbol, Ensembl gene ID, transcript ID, or hg38 genomic coordinates. Coordinate-based queries retrieve nucleosomes overlapping a user-specified genomic interval, with a maximum query span of 5 Mb. For each selected nucleosome, OncoNucAtlas provides its genomic annotation together with an interactive visualization of nucleosome occupancy across samples and cancer types. For cancer-type-level nucleosomes, mean occupancy with the standard error of the mean was displayed for the selected cancer type. For cancer-type-enriched nucleosomes, sample-level occupancy distributions are displayed as box plots to facilitate comparison between the cancer type to which the nucleosome was assigned and user-selected reference cancer types.

## Author contributions

Y.L. conceived and supervised this project. S.Y. performed most of the analysis. and J.Y. and J. Y performed part of the analysis. Y.L. wrote most part of the manuscript and S.Y. wrote some part. All authors interpreted the results, read, revised, and approved the final manuscript.

## Acknowledgments

The authors thank members of Yumei Li’s lab and Wei Li’s lab for helpful discussions.

## Supplementary information

Supplementary Figs. S1 to 12, and Tables S1 to 16.

