## Supplementary figures for "Pan-cancer nucleosome landscapes reveal tumor heterogeneity and inform cell-free DNA-based detection"

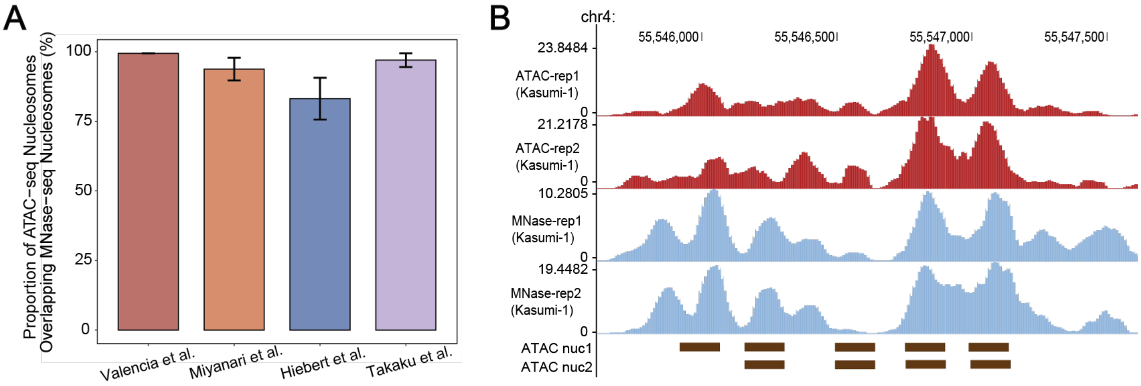


**Fig. S1. ATAC-seq-derived nucleosome signals recapitulate MNase-seq-defined nucleosome patterns.**

**(A)** Proportions of ATAC-seq-inferred nucleosomes overlapping with MNase-seq-defined nucleosomes across four cell-line datasets (from left to right: TTC1240, eHAP, Kasumi-1, and MDA-MB-231). Bars represent the mean across samples, with error bars indicating standard deviation (SD).

**(B)** Representative nucleosome occupancy profiles derived from ATAC-seq (red) and MNase-seq (blue) in the Kasumi-1 cell line. ATAC-seq-inferred nucleosome positions are shown below the occupancy tracks.

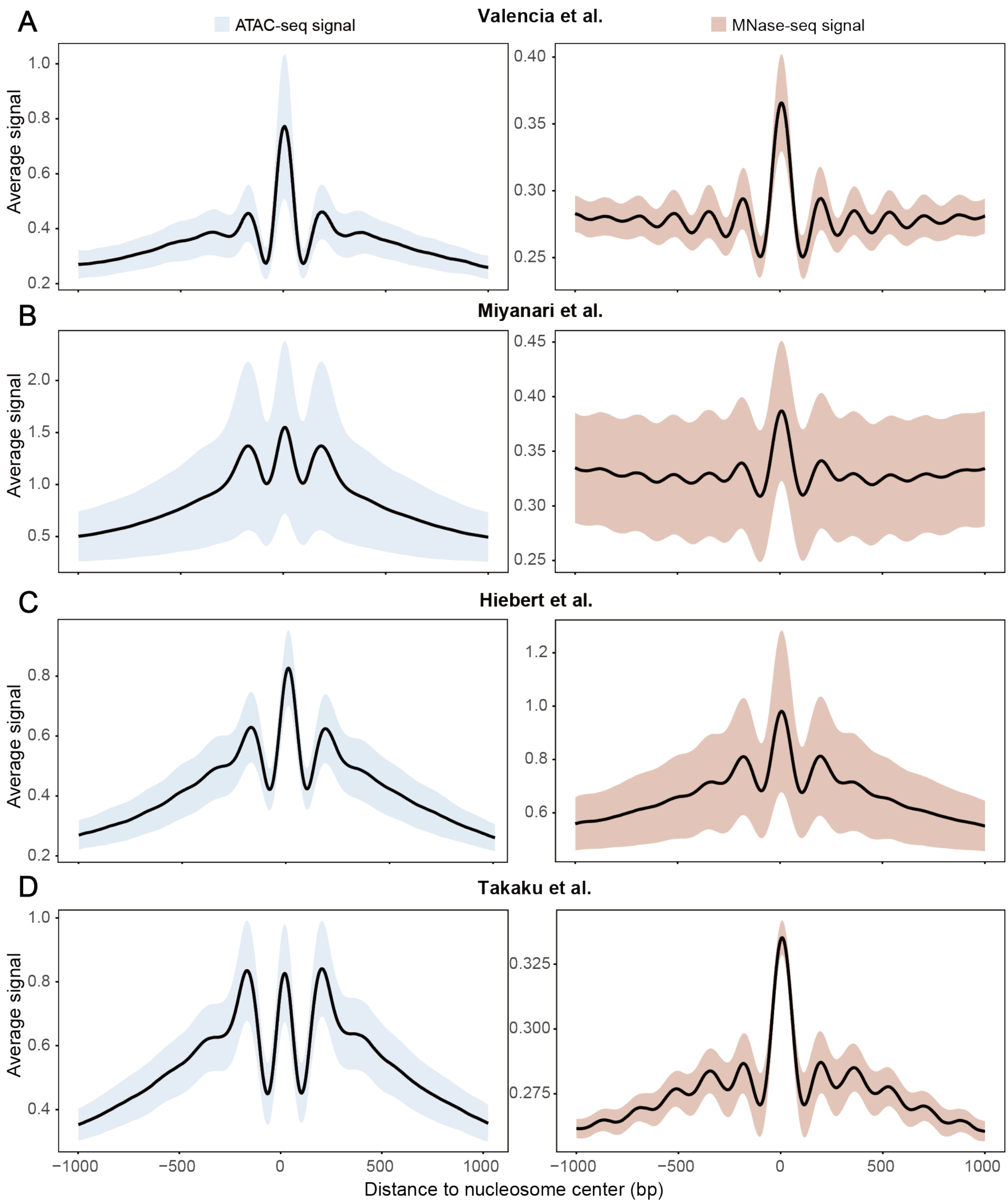


**Fig. S2. ATAC-seq-derived nucleosome positions capture characteristic MNase-seq occupancy patterns.** Aggregative ATAC-seq and MNase-seq nucleosome occupancy profiles around ATAC-seq-inferred nucleosome positions in TTC1240 **(A)**, eHAP **(B)**, Kasumi-1 **(C)**, and MDA-MB-231 **(D)**. Lines represent the mean occupancy profiles, with shaded areas representing 95% confidence intervals.

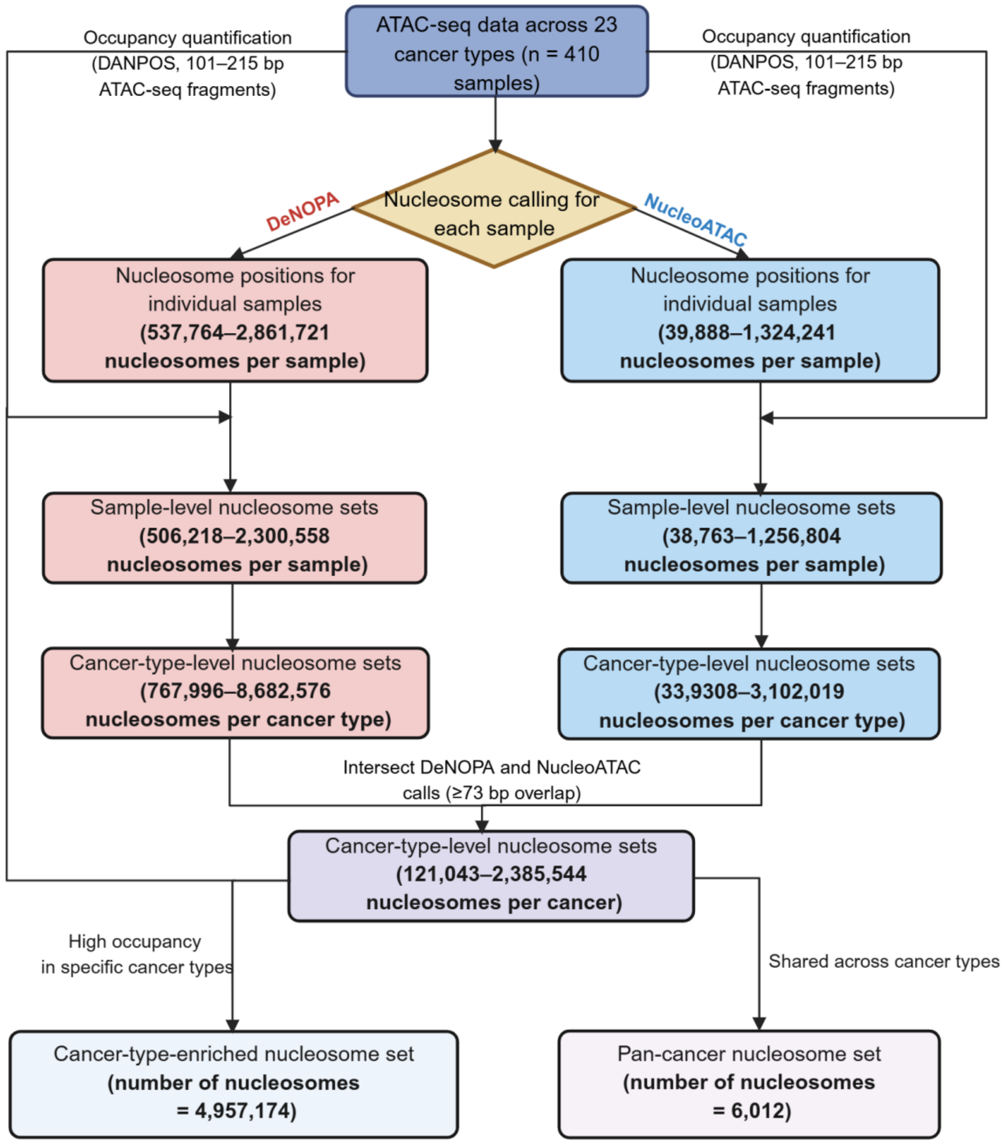


**Fig. S3. Workflow for constructing the pan-cancer nucleosome atlas and identifying cancer-type-enriched nucleosomes.** ATAC-seq data from 410 primary tumors across 23 cancer types were used to infer sample-level nucleosome positions. Reproducible nucleosomes were integrated to generate cancer-type-level nucleosome sets, from which pan-cancer and cancer-type-enriched nucleosomes were identified.

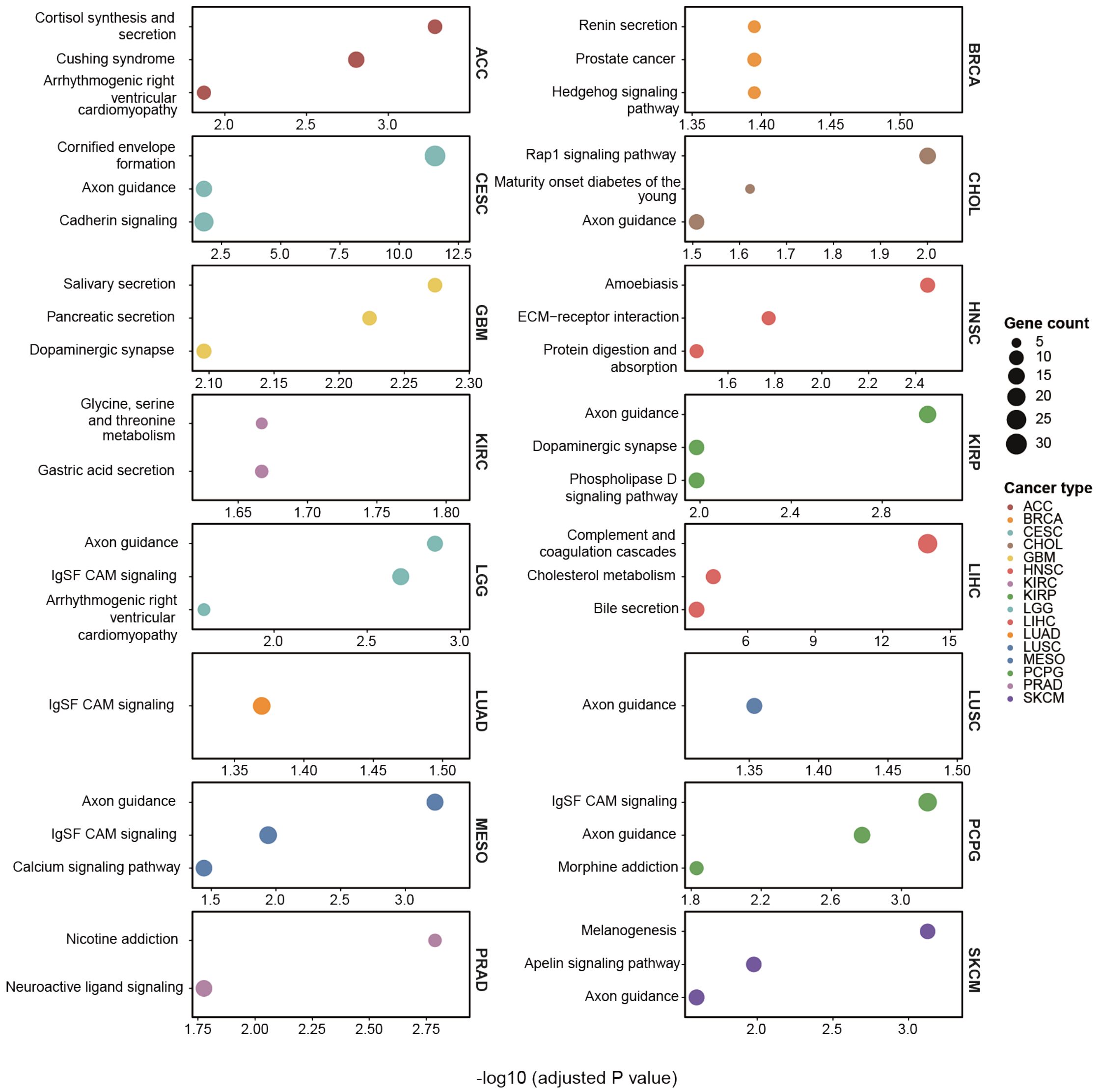


**Fig. S4. Cancer-type-enriched nucleosomes reveal cancer-specific biological pathways.** KEGG pathway enrichment analysis of genes associated with cancer-type-enriched nucleosomes for each cancer type. The top three significantly enriched pathways ranked by adjusted P value are shown for each cancer type. Bubble size represents gene count, and the x-axis represents −log10(adjusted P value).

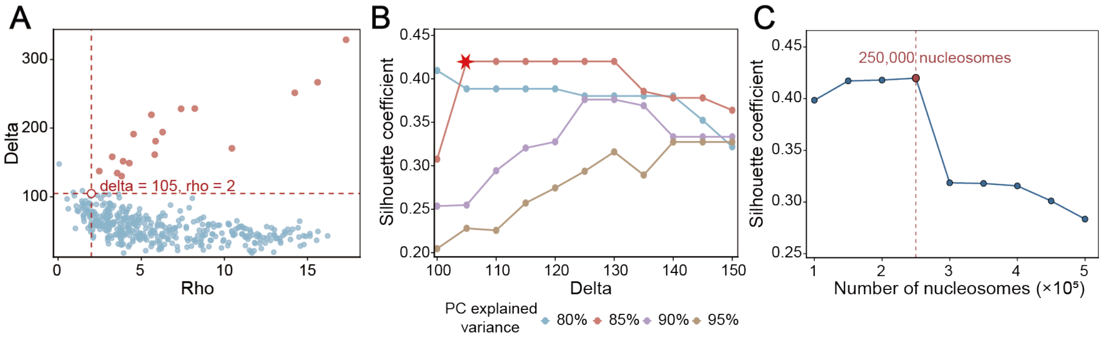


**Fig. S5. Optimal nucleosome-based clustering with the selected parameter combination.**

**(A)** Decision plot for density-based clustering using the 250,000 most variable cancer-type-enriched nucleosomes and principal components explaining 85% of the cumulative variance.

**(B)** Mean silhouette coefficients across different PC cumulative variance thresholds and delta values. The asterisk indicates the parameter combination with the highest mean silhouette coefficient.

**(C)** Mean silhouette coefficients across different numbers of the most variable cancer-type-enriched nucleosomes, with the PC cumulative variance threshold fixed at 85% and delta fixed at 105.

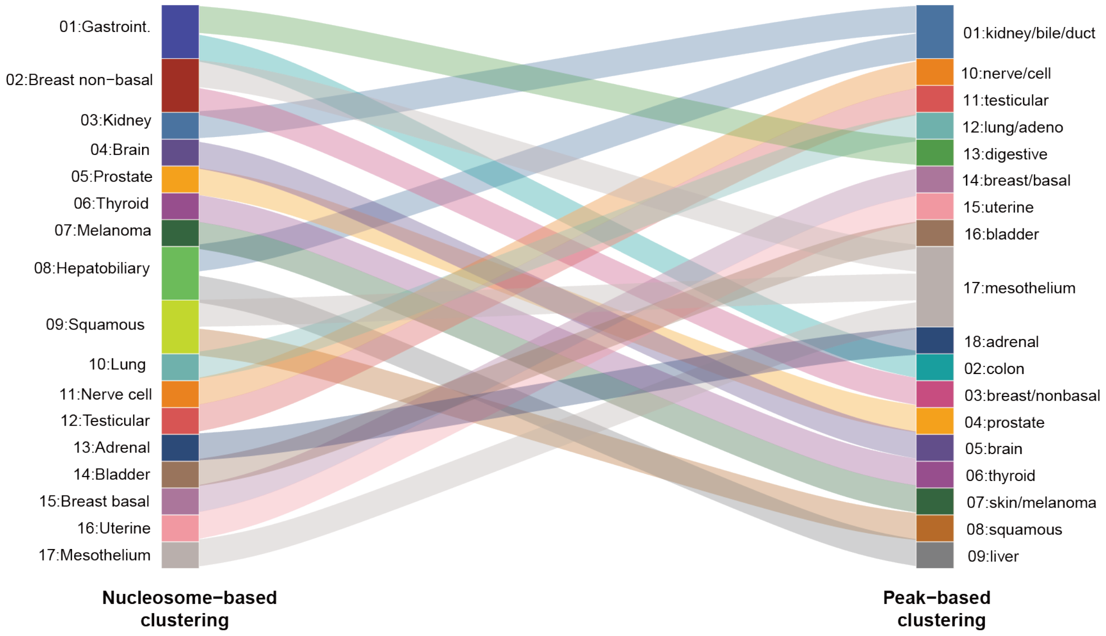


**Fig. S6. Nucleosome positioning reveals additional tumor structure beyond chromatin accessibility.** Alluvial plot showing the correspondence between the clusters identified from cancer-type-enriched nucleosome occupancy profiles and cancer-type-enriched ATAC-seq peak profiles. Flows connect corresponding clusters between the two clustering approaches.

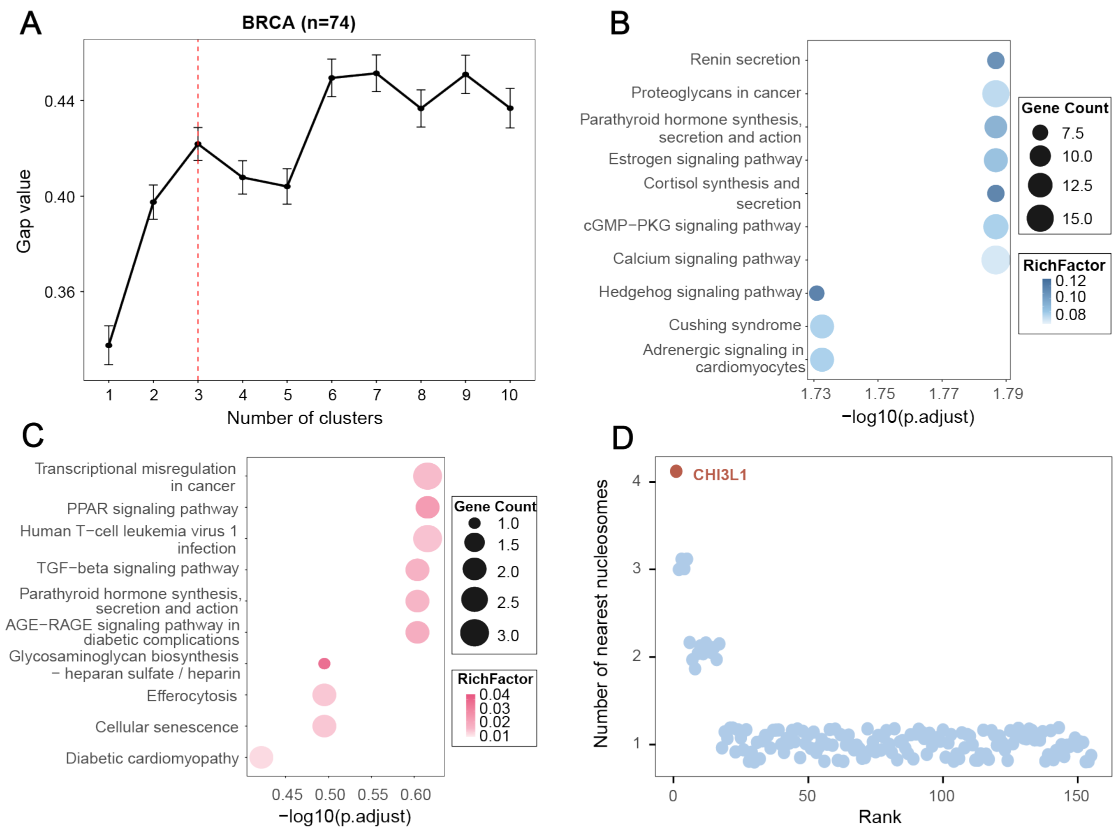


**Fig. S7. Nucleosome-defined BRCA subgroups exhibit distinct molecular features.**

**(A)** Gap statistic analysis for determining the optimal number of nucleosome-defined BRCA subgroups.

**(B–C)** KEGG pathway enrichment of genes associated with subgroup 2-specific **(B)** and subgroup 3-specific **(C)** nucleosomes. Bubble size represents gene count, bubble color represents the rich factor, and the x-axis represents enrichment significance as −log10(adjusted P value).

**(D)** Ranking of genes by the number of subgroup 1-specific promoter-associated nucleosomes, highlighting CHI3L1.

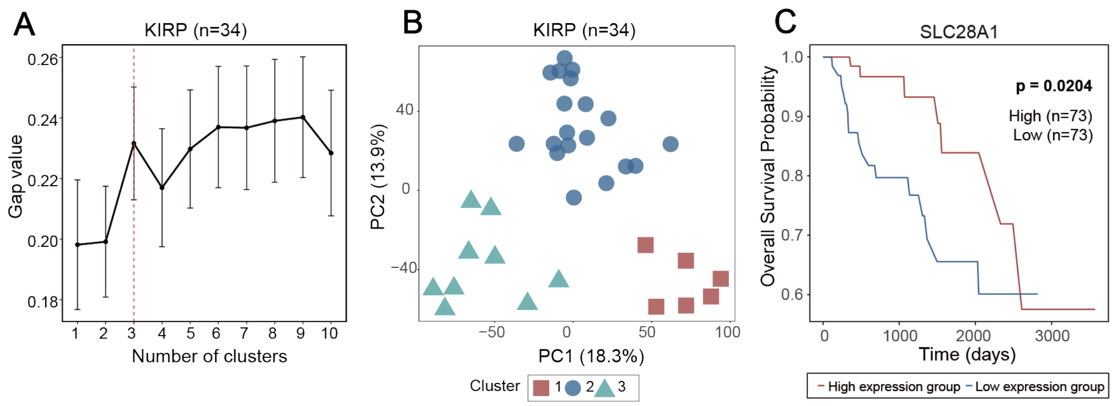


**Fig. S8. Nucleosome occupancy patterns reveal molecular heterogeneity among KIRP subgroups.**

**(A)** Gap statistic analysis for determining the optimal number of nucleosome-defined KIRP subgroups.

**(B)** Principal component analysis (PCA) of KIRP samples based on nucleosome occupancy profiles, colored by nucleosome-defined subgroups.

**(C)** Subgroup 1-specific promoter-associated nucleosome alterations linked to SLC28A1 and Kaplan–Meier survival analysis comparing KIRP patients with high and low SLC28A1 expression.

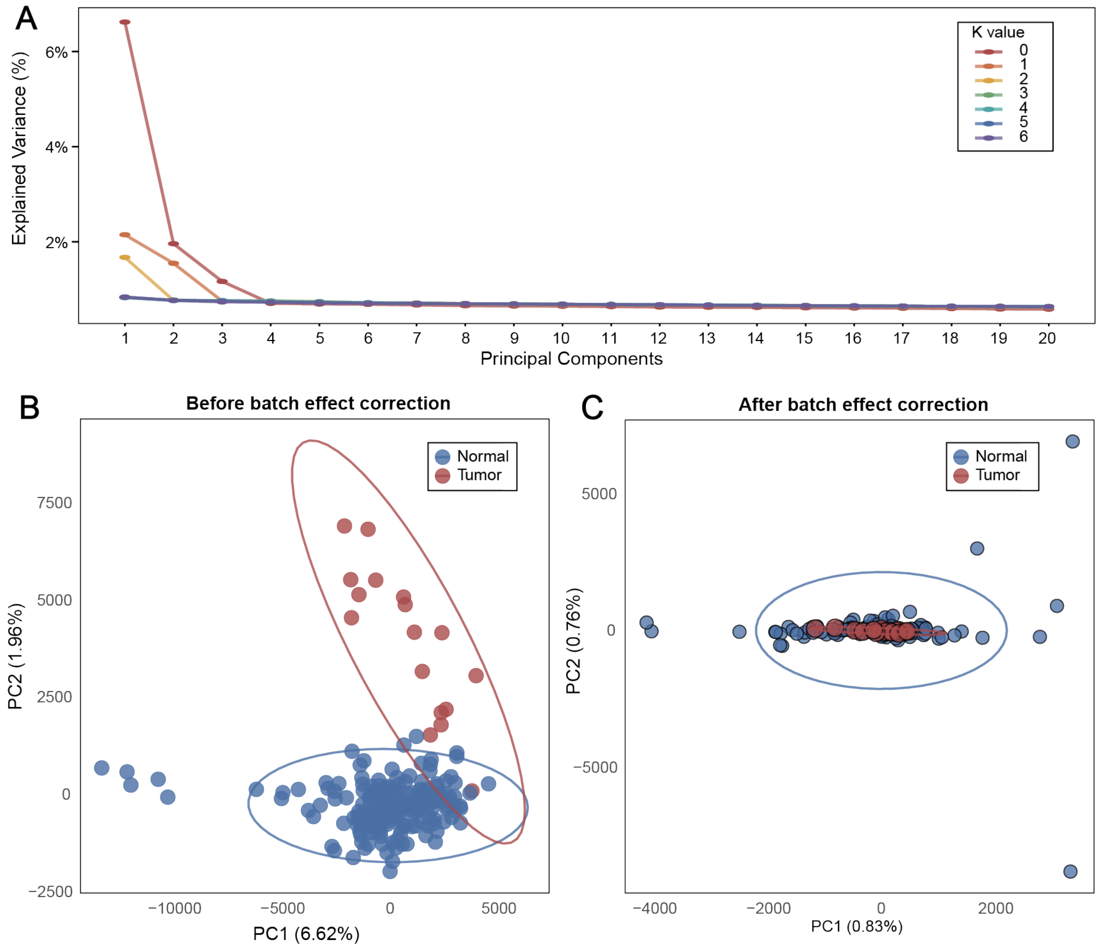


**Fig. S9. RUVg correction reduces batch-associated variation in tumor and normal liver samples.**

**(A)** Variance explained by the first 20 principal components before RUVg correction (k = 0) and after correction with k values from 1 to 6.

**(B–C)** PCA of liver tumor and normal samples before **(B)** and after **(C)** RUVg batch-effect correction (k = 4).

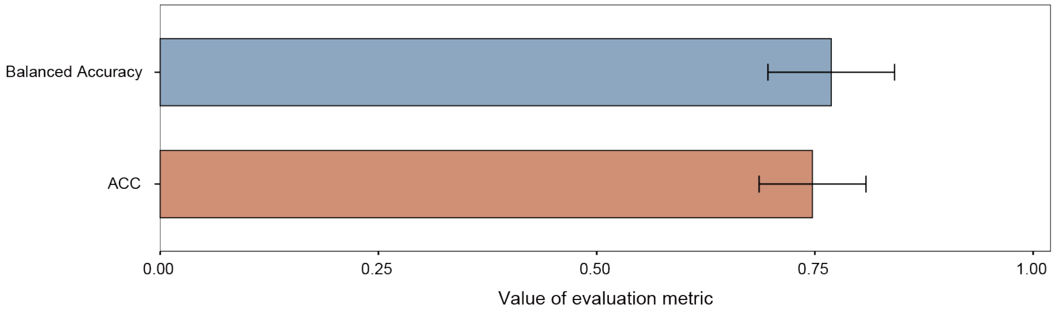


**Fig. S10. Cancer-type-enriched nucleosomes enable multiclass cancer classification.** Balanced accuracy and accuracy (ACC) of the cancer multiclass classifier evaluated by 10-fold cross-validation. Bars represent the mean across folds, with error bars indicating standard deviation (SD).

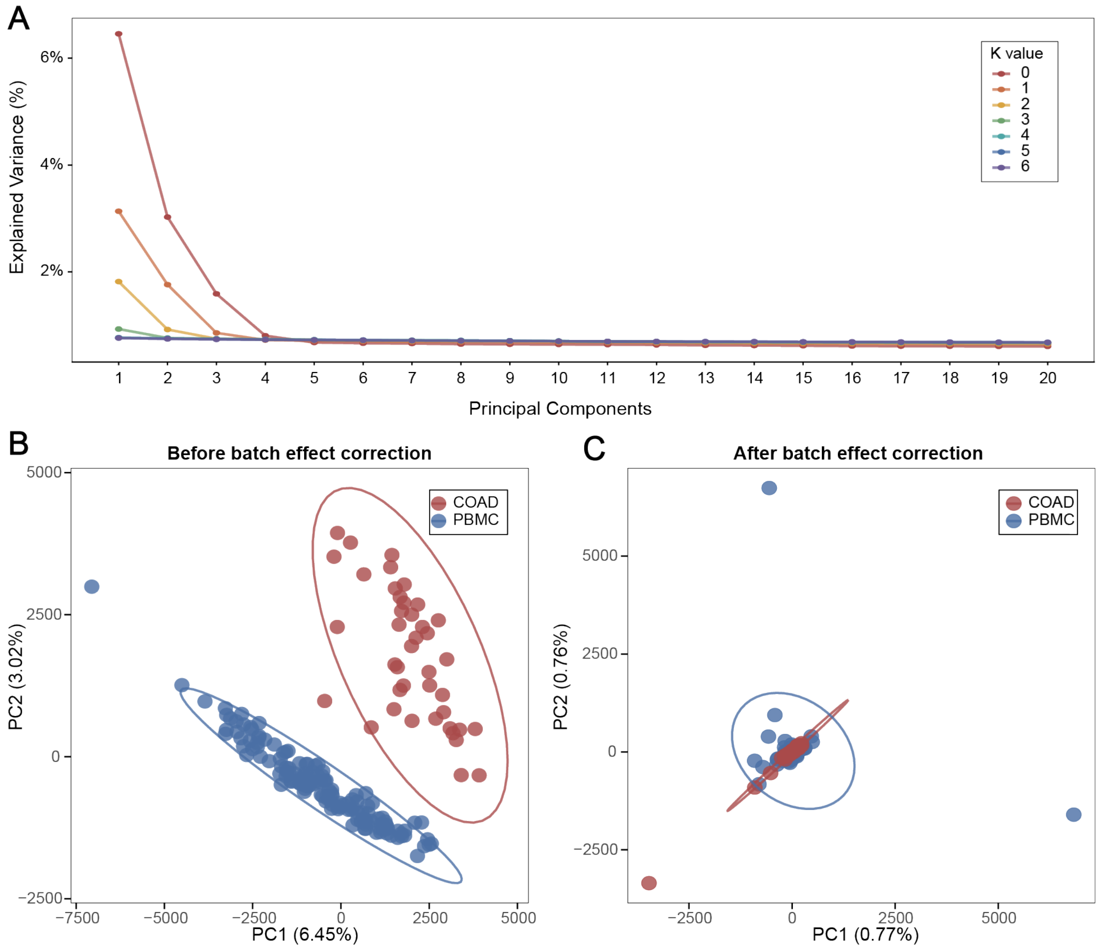


**Fig. S11. RUVg correction reduces batch-associated variation in tumor tissues and PBMCs.**

**(A)** Variance explained by the first 20 principal components before RUVg correction (k = 0) and after correction with k values from 1 to 6 for COAD.

**(B–C)** PCA of COAD tumor and PBMC samples before **(B)** and after **(C)** RUVg batch-effect correction (k = 4).

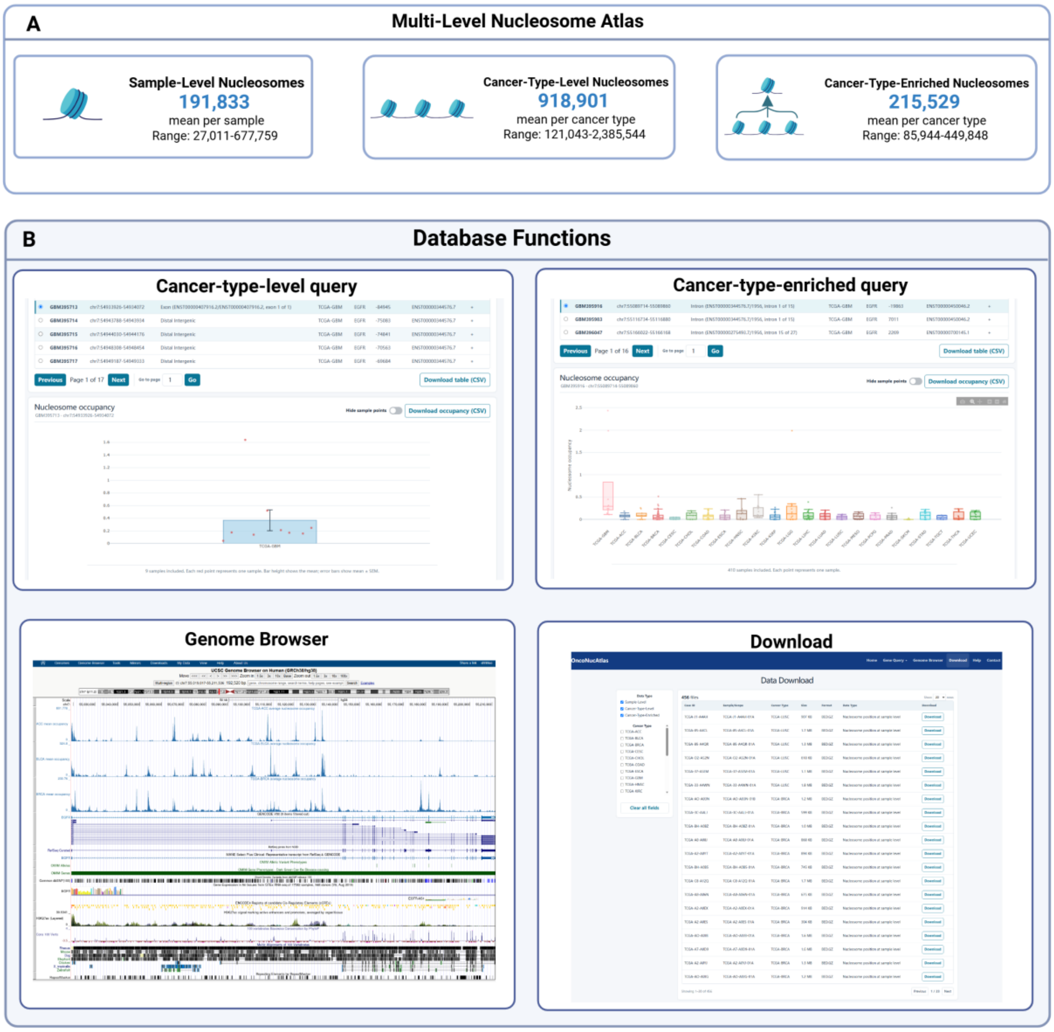


**Fig. S12. OncoNucAtlas provides a comprehensive resource for exploring cancer nucleosome landscapes.**

**(A)** Overview of sample-level, cancer-type-level, and cancer-type-enriched nucleosome datasets included in OncoNucAtlas.

**(B)** Major functions of the OncoNucAtlas web platform, including nucleosome querying, cross-cancer occupancy comparison, data download, and UCSC Genome Browser visualization.
